# A Chromosome-Level Assembly of the Pine Processionary Moth (*Thaumetopoea pityocampa*) genome

**DOI:** 10.1101/2025.06.06.658235

**Authors:** Mathieu Gautier, Charles Perrier, Pierre Nouhaud, Jacques Lagnel, Manuela Branco, Thomas Chertemps, Franck Dorkeld, Marie-Christine François, Bernhard Gschloessl, Frédérique Hilliou, Emmanuelle Jacquin-Joly, Fabrice Legeai, Gaëlle Le Goff, Céline Lopez-Roques, Martine Maibeche, William Marande, Hugues Parinello, Laure Sauné, Charles Perrier, Carole Kerdelhué

**Affiliations:** CBGP, INRAE, CIRAD, IRD, Institut Agro, Univ. Montpellier, 755 avenue du Campus Agropolis, CS30016, F-34988 Montferrier-sur-Lez Cedex, France; GAFL, INRAE, Domaine Saint Maurice, Allée des Chênes, 84140 Montfavet, France; Centro de Estudos Florestais (CEF), Instituto Superior de Agronomia (ISA), University of Lisbon, Lisbon, Portugal; Centro de Estudos Florestais (CEF), Laboratório Associado TERRA, Instituto Superior de Agronomia (ISA), University of Lisbon, Lisbon, Portugal; Institut d’Ecologie et des Sciences de l’Environnement de Paris (iEES-Paris), Sorbonne Université, CNRS, INRAE, IRD, Université Paris Cité, Université Paris Est Créteil, F-75005 Paris, France; INRAE, Sorbonne Université, CNRS, IRD, Université Paris Cité, Université Paris Est Créteil Val de Marne, Institut d’Ecologie et des Sciences de l’Environnement de Paris (iEES-Paris), 78026 Versailles cedex, France; Université Côte d’Azur, INRAE, CNRS, ISA, F-06903 Sophia Antipolis, FRANCE; IGEPP, BioInformatics Platform for Agroecosystems Arthropods (BIPAA), Rennes, France; Univ Rennes, Inria, CNRS, IRISA, 35000, Rennes, France; INRAE, GeT-PlaGe, Genotoul, 31326 Castanet-Tolosan, France; CNRGV, INRAE, Castanet Tolosan, France; MGX-Montpellier GenomiX, Univ Montpellier, CNRS, INSERM, Montpellier France

**Keywords:** *Thaumetopoea pityocampa*, Pine Processionary Moth, Genome Assembly, ONT

## Abstract

We present a chromosome-level genome assembly and annotation of the pine processionary moth, *Thaumetopoea pityocampa* (Lepidoptera: Notodontidae), a key forest pest that is a public health concern. The nuclear genome spans 615.9 Mb, scaffolded into 50 chromosome scale and 115 smaller scaffolds, with high completeness (BUSCO score: 98.9%) that provides a decisive improvement over the previous assembly (537 Mb; 68,292 contigs; BUSCO 83.6%). Coverage differences in resequenced males and females allowed identification of the Z chromosome and several W-linked contigs. As expected from previous studies, we found that synteny was largely conserved with related Lepidoptera, although chromosomal fissions may explain the higher chromosome number of 49 autosomes compared to typical lepidopteran karyotypes. We also integrated into the assembly linkage map, allowing estimation of a genome-wide male recombination rate of 5.06 cM/Mb, varying from 11.6 cM/Mb to 1.98 cM/Mb from the smallest to the largest chromosomes. Repetitive elements represented 49.1% of this new assembly, dominated by LINEs (45.1% of classified repeats). Finally, gene prediction identified 12,898 gene models, among which 17 circadian rhythm genes were manually curated. Expert annotation further allowed to identify 51 genes of the odorant receptor (OR) family as well as a total of 236 detoxification genes, including 78 CYPs, 56 CCEs, 30 GSTs, 23 UGTs and 49 ABCs. Overall, this assembly represents the first chromosome-level genome for a member of the Thaumetopoeinae subfamily, significantly expanding the currently limited set of genomic resources available for Notodontidae. The fully annotated assembly is publicly accessible through the LepidoDB database (https://bipaa.genouest.org/is/lepidodb/) and will serve as a valuable resource for research on population genomics of this species.

## Introduction

The pine processionary moth (PPM), *Thaumetopoea pityocampa* (Dennis & Schiffermüller; Lepidoptera: Notodontidae), is one of the main defoliators of pines in southern Europe and North Africa. It is also a health concern for humans and animals due to the urticating setae carried by its late larval instars (Battisti *et al*., 2017). The species is univoltine throughout its distribution range (except during prolonged diapause), with mating and oviposition in summer and gregarious larval development spanning autumn and winter.

*T. pityocampa* is responding to climate change through northward range expansion in Europe (Rossi *et al*., 2025) and local declines at its southern edge in North Africa (Bourougaaoui *et al*., 2021). Geographic variation in phenology has been observed, suggesting local adaptation (Robinet *et al*., 2015; Martin *et al*., 2022), including an allochronic population in Portugal (the so-called Summer Population or SP) with a shifted reproductive period that reduces gene flow and can lead to incipient speciation (Pimentel *et al*., 2006; Burban *et al*., 2016; Santos *et al*., 2011). These characteristics make *T. pityocampa* an excellent model for studying the genomics of range expansion, adaptive response to climate change, and allochronic divergence.

Previous genomic resources included a draft genome and transcriptome focused on the aforementioned Portuguese allochronic population (Gschloessl *et al*., 2018), but these were highly fragmented (almost 70,000 scaffolds) and did not meet current standards for population or comparative genomics. In this study, we present a new high quality chromosome-level genome assembly for *T. pityocampa* based on Oxford Nanopore Technologies (ONT) long-read sequencing and Hi-C scaffolding.

In addition to the improved assembly, our work integrates genetic linkage map data and provides a comprehensive functional and structural annotation of the genome, including transposable element (TE) content and gene models. We also provide expert annotation of genes involved in two important functions in insects, namely olfaction and detoxification. The first group allows species to use volatile signals to locate food sources, mating partners, oviposition sites, and to avoid danger (Leal, 2013). The second is involved in the metabolism of both exogenous compounds, such as insecticides or allelochemicals emitted by host plant, and endogenous compounds, such as hormones and vitamins (Després *et al*., 2007) These genomic resources establish a critical foundation for future research on the evolutionary ecology and genetics of this species.

## Methods and Materials

### Construction of a contig assembly using ONT long reads

L5 larvae were collected alive from a single nest in a black pine (*Pinus nigra*) in February 2018 in Orléans La Source, France. The larvae were immediately flash frozen in liquid nitrogen and stored at −80°C until DNA extraction. High molecular weight (HMW) DNA was extracted from two larvae using an in-house protocol. The extracted DNA was then sent to the GenoToul platform, and used to prepare an ONT library using the Ligation Sequencing Kit (SQK-LSK109, Oxford Nanopore Technologies) following the manufacturer’s instructions, specifically the “1D gDNA selecting for long reads” protocol. At each step, DNA quantity was measured using the Qubit dsDNA HS Assay Kit (Life Technologies), and purity was assessed using a Nanodrop spectrophotometer (Thermo Fisher Scientific). Size distribution and potential degradation were evaluated using the Fragment Analyzer (Agilent) with the DNF-464 HS Large Fragment Kit. Purification steps were performed using AMPure XP beads (Beckman Coulter). A total of 7 µg of purified DNA was obtained, which was subjected to an additional DNA repair step using the SMRTbell SPV3 damage repair kit (PacBio). To ensure the integrity of long reads, a final size selection step was performed using the Short Read Eliminator XS Kit (Circulomics) without prior shearing. The size-selected DNA (2.5 µg) was then used to prepare the sequencing library with the SQK-LSK109 kit (ONT).

The library was initially tested on an R9.4.1 Flongle flow cell and sequenced for 24 hours on a GridION instrument at 8 fmol. Subsequently, it was loaded onto an R9.4.1 flow cell and sequenced for 72 hours on a PromethION instrument at 16 fmol. Basecalling was performed using MinKNOW-Live-Basecalling (ONT, UK), versions 3.6.0 and 3.6.1 for the respective runs, with default settings yielding a total of 39,972 reads (513 Mb, N50=21,316 bp) and 6,945,599 reads (102.4 Gb, N50=22,070 bp) as detailed in Table S1. The reads were then trimmed using `Porechop` (v0.2.3; Wick *et al*., 2017) ran with default options, combined and self-corrected using the `Canu` assembler (v1.9; Koren *et al*., 2017) with options `-correct saveReadCorrections=true` (to run only the error correction step); `minReadLength=1000` (to discard reads shorter than 1,000 bp); `stopOnReadQuality=false` (to pro-cess all reads regardless of quality); `genomeSize=540m` (setting an assumed genome size of 540 Mb as in Gschloessl *et al*., 2018); and `corOutCoverage=100` (to use up to 100X coverage for correction). After the correction step, a total of 1,874,661 high-quality reads (53.67 Gb, N50=29,206 bp) were available for genome assembly which was performed using `SMARTdenovo` (v.1.0.0.; Liu *et al*., 2021) run with default settings, and which resulted in an assembly consisting of 362 contigs totaling 632,816,461 bp with an N50=5,549,426 (and N90=1,339,262).

This assembly was further polished using Whole-Genome short-read sequences generated for the same DNA sample (i.e., the two L5 larvae) used for the ONT library. To that end, two DNA-seq libraries were prepared using the Illumina TruSeq Nano DNA HT Library Prep Kit (Illumina, California, USA). Briefly, 300 ng of DNA was fragmented to ca. 550 bp using an M220 Focused-Ultrasonicator (COVARIS). Size selection was performed with SPB beads, and adapters were ligated for sequencing. Library quality was assessed using an Agilent Fragment Analyzer with the DNF-474-0500 High Sensitivity NGS Kit, and library quantification was performed via qPCR using the Kapa Library Quantification Kit (Roche). The library was then sequenced on an Illumina HiSeq 3000 with a paired-end (PE) read length of 2×150 bp, using the HiSeq 3000 Reagent Kits. A total of 350.6 millions read pairs were obtained (totaling 107.8 Gb, i.e. 170X coverage assuming a genome size of 632.8 Mb); and filtered using fastp (v0.20.0; Chen *et al*., 2018a) run with options `--detect_adapter_for_pe` and `--average_qual 25 --low_complexity_filter --complexity_threshold 30`. The filtered reads (342.6 millions pairs totaling 102.7 Gb) were then used to polish the contig assembly with two consecutive runs of Pilon (v1.23; Walker *et al*., 2014) run with default options.

The resulting polished assembly consisted of 362 contigs totaling 636.2 Mb (N50=5.58 Mb). The overall %GC was equal to 37.61 consistent with that observed on raw individual WGS data. Yet, the size of the assembly remained significantly larger than the previous assembly (GS=537 Mb, Gschloessl *et al*., 2018) and than the haploid genome size (from 564 to 573 Mb, Figure S1) we esti-mated with `GenomeScope` from the k-mer spectrum computed with `JellyFish` (Marçais and Kings-ford, 2011) from the short-read WGS data we generated for two males and two females primarily to discriminate autosomal and sex-chromosome contigs (see below and Table S2). This suggested some contamination of our assembly by exogenous sequence (e.g., bacterial endosymbionts) which was also supported by the %GC of the different contigs (Figure S2). The primary assembly was thus filtered by removing 23 contaminant contigs (i.e., non lepidopteran) identified following Gautier (2023) by querying a database constructed from the NCBI non-redundant nucleotides (nt) released in February 2020 using the program `Kraken2` (v2.1.2; Wood *et al*., 2019). These included three large contigs with %GC*>* 50 (of 3.53 Mb, 2.16 Mb and 0.813 Mb) sharing similarity with bacteria of the *Pseudomonas* genus. In addition, two mitochondrial contigs of length 89.6 kb (%GC=34.6) and 70.6 kb (%GC=37.0) were identified by aligning the CO1 mitochondrial gene sequence for *T. pityocampa* (NCBI: MH742439), and found to align to other lepidopteran mitochondrial genomes over large stretches of sequences (up to 6 kb) and also to each other over more than 10 kb. As the mitochondrial genome is circular, it is actually expected that our assembly approach may not recover a complete and accurate mitochondrial sequence. We thus discarded these two mitochondrial contigs and replaced them by a *de novo* mitochondrial sequence assembled from the corrected ONT reads using the `MitoFinder` program (v1.4.1; Allio *et al*., 2020). Last, a refined search for alternative chromosomal copies using the `PurgeHaplotigs` pipeline (Roach *et al*., 2018) was used to flag and remove a total of 105 additional problematic sequences (repeat-rich contigs or haplotigs) from the assembly.

### HiC scaffolding

To further scaffold the contig assembly at the chromosome level, we incorporated HiC data, which provide long-range chromatin interaction information. The HiC library was prepared in the MGX facility using the Arima-HiC kit (according to the manufacturer’s instructions) from a pool of ten flash frozen L2 caterpillars that were collected in September 2020 in a black pine in the same locality (Orléans La Source, France) as in 2018, kept alive and fed with black pine needles for a few days. The HiC library was then sequenced on an Illumina NovaSeq 6000 to generate 284,064,351 pairs of 150 bp reads. These raw paired-end reads were then `filtered` using fastp (v0.19.4; Chen *et al*., 2018b) with default options, leading to 565,166,160 available sequences (84.72 Gb, i.e., *>*130X genome coverage) with an estimated 21.6% of duplicate reads. The filtered paired-end reads were then mapped to the filtered contig assembly using the HiCstuff pipeline program (v3.0.2; Baudry *et al*., 2020; Matthey-Doret *et al*., 2020) with the options `--mapping=iterative, --read-len=150, --duplicates, --distance-law, --filter` and `-e “DpnII,HinfI”`. A total of 115,468,787 valid pairs (i.e., 40.90% of the initial pairs) were successfully identified and used to scaffold the assembly with `instaGRAAL` (v0.1.6; Baudry *et al*., 2020) run with options -l 6 (i.e., contact map was binned into sets of 3^6^ = 729 consecutive restriction fragments leading to a total of 3,470 bins on the contig assembly) and -n 50 (i.e., 50 MCMC cycles ensuring convergence of the algorithm as empirically confirmed by examining the likelihood trace). The scaffold assembly obtained was finally polished to favor the initial orientation of the fragments within the contig (Baudry *et al*., 2020) using a slightly modified version of the `instagraal-polish` program. As a first empirical quality assessment of the resulting scaffolded assembly (named `Tpit_2.0` hereafter), we finally aligned the sequences of 11 Bacterial Artificial Chromosomes (BACs) (Gschloessl *et al*., 2018), ranging from 40 to 128 kb using `dgenies` (v1.5.0; Cabanettes and Klopp, 2018) run with default options. Although the BAC library was constructed from samples of a distantly related Portuguese population, all BACs showed high similarity and contiguous alignments with only one scaffold, with a mean overall nucleotide divergence of 0.657% over a cumulative length of 673,859 bp from alignment blocks *>* 10*kb* (Figure S3). Yet, some preliminary analyses focusing on specific regions of the fifth-largest scaffold (referred to hereafter as chromosome 4) motivated us to fill the gap between two consecutive contigs. To that end, we identified and sequenced a BAC (B29F14) overlapping this region, using an approach similar to that described in Gschloessl *et al*. (2018). The resulting BAC sequence, consisting of a single contig of 61,721 bp, was then used to manually fill the gap and resolve small inconsistencies at its boundaries. The final updated scaffold assembly is hereafter referred to as Tpit 2.1. A snailplot was produced using the `BlobToolKit` viewer (v4.3.5; Challis *et al*., 2020) to visualize key assembly metrics. Assembly completeness was assessed with BUSCO v5.7.1 using the `lepidoptera_odb10` reference dataset, which contains 5,286 conserved genes (Manni *et al*., 2021). Finally, we searched for telomeric sequence footprints at the ends of the scaffolds using the find module from the tidk toolkit (v0.2.65; Brown *et al*., 2025), with the option `-c Lepidoptera` to count occurrences of the *AACCT* motif, commonly found in the telomeres of Lepidoptera species, in nonoverlapping 10 kb windows.

### Identification of sex-linked and autosomal scaffolds

Sex-linked and autosomal scaffolds were identified, following the methodology described in Gautier *et al*. (2018), by comparing the overall read coverage of their underlying contigs, using short-read sequencing data from two males and two females emerged in laboratory conditions from pupae sampled in July 2012 in the summer population (SP) from the Mata Nacional de Leiria (Portugal). Briefly, DNA paired-end libraries with insert size of ca. 350 bp were prepared using the Illumina TruSeq Nano DNA Library Preparation Kit following manufacturer protocols for each individual identified with a specific barcode. Libraries were then validated on a DNA1000 chip on a Bioanalyzer (Agilent) to determine size and quantified by qPCR using the Kapa library quantification kit to determine concentration. The cluster generation process was performed on cBot (Illumina Inc.) using the Illumina Paired-End DNA sample preparation kit (FC-102-1001, Illumina Inc.). Each individual library was further paired-end sequenced (2×125 bp) on an Illumina HiSeq 2500 instrument using the Sequence by Synthesis technique. Raw paired-end reads were then filtered using fastp (v0.19.4; Chen *et al*., 2018b) run with default options to remove contaminant adapter sequences and trim for poor quality bases. Details on the four Ind-Seq data sets are given in Table S2. The filtered reads were then aligned onto the contig assembly using default options of the `mem` program from the `bwa` software package (v0.7.12; Li, 2013). Read alignments with a MAQ*<*20 and PCR duplicates were further removed using the `view` (option -q 20) and `rmdup` programs from the `samtools` software (v1.9; Li *et al*., 2009). As shown in Table S2, the total number of mapped reads per individual after filtering ranged from 132,621,113 to 154,281,790 reads (with a proportion of properly paired reads ranging from 97.3% to 97.5%) resulting in a high realized coverage (ranging from 29X to 33X). Read coverage at each position of the assembly for the four individual `bam` files was then jointly estimated using the `depth` program from the `samtools` software (v1.9; Li *et al*., 2009) run with default option. To limit redundancy, only one count every 100 successive genomic positions were retained for further analysis.

Following Gautier *et al*. (2018), we estimated for each and every contig the ratio, denoted *ρ* hereafter, of the relative (average) read coverage of contigs between ZZ males and ZW females. The ratio *ρ* is expected to be equal to 1 for autosomal contigs, to 2 for Z chromosome contigs and to 0 for W chromosome contigs. More precisely, let *c_ijk_* represent the observed coverage in individual *k* for the contig *i* at the *j*^th^ position among the 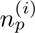 covered. Note that 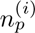 should be approximately equal to *s_i_/*100 where *s_i_* is the contig size in bp. We then define female (male) read coverage 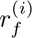 (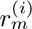) for contig *i* as the average across the 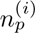 positions read coverages over all the *n_f_* = 2 females (*n_m_* = 2 males): 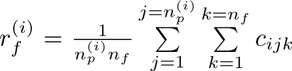 and 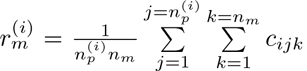. To account for possible differences in the realized sequencing coverage for males and females, the ratio *ρ_i_* was then estimated for each contig *i* as: 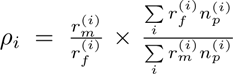. Visual inspection of the estimates suggested that the ratio were inaccurate (outlying values) for the small contigs and/or displaying extreme GC contents. As a matter of expedience, we thus chose to discard contigs *<* 150 kb in length and/or with a %GC*>* 39.2 (third distribution quartile), as well as low covered contigs (*<*5X in all four individuals). Overall, 199 contigs (out of the 232) totaling 612.5 Mb (out of 615.7 Mb of the contig assembly) were retained for further analysis. We then fitted a Gaussian mixture model with three classes with unknown means and variances on the 199 estimates of *ρ* to classify the underlying in three classes corresponding to autosomal, Z– and W–linked contigs. The latter parameters were estimated using an Expectation-Maximization algorithm as implemented in the `mixtools` package (v1.1.0 Benaglia *et al*., 2009) for the R software (R Core Team, 2017). This leads to the delineation of three distinct clusters with estimated means 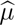_*W*_ = 0.117 ± 0.077, 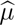_*A*_ = 0.971±0.023 and 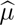_*Z*_ = 1.92±0.055 that were interpreted as representing W–linked, autosomal and Z–linked contigs, respectively. As expected, the estimated average ratio for the autosomal and Z–linked classes was lower than 1 and 2, respectively, due to the inclusion of Z–linked contigs in the overall normalizing denominator of *ρ* (as confirmed *a posteriori* after re-estimating considering only contigs assigned to autosomes for normalization). Conversely, the mean of the first class was non-null (but displayed the largest estimated variance), likely due to the presence of non-specific W sequences in the genome making estimated coverages of males non-null for this chromosome. To classify the different contigs we finally computed (two-sided) p–values for each contig associated with each chromosome type *K* (i.e., *K* = *Z*, *K* = *A* and *K* = *W*, for the Z, autosome and W respectively) as: 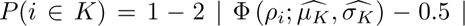 (where Φ(*x*) represents the Gaussian cumulative distribution function). Contigs were assigned to a chromosome type *K* if and only if *P* (*i* ∈ *K*) *>* 0.01. This leads to the unambigous assignation of 173 autosomal (totaling 574.6 Mb), 4 Z–linked (totalling 28.97 Mb) and 17 W–linked (totalling 4.909 Mb) contigs; while only 15 contigs (totaling 4.073 Mb) out of the 199 investigated contigs were left unassigned.

As a final quality check we compared the read coverage of the different contigs as a function of their assigned chromosome types for sequencing data obtained on the DNA pool of the two larvae (of unknown sex) sampled in Orléans that were used to build the assembly. To that end, the short-reads generated to polish the assembly were filtered using `fastp` (v0.23.4; Chen *et al*., 2018a) run with default options and then mapped onto the (cleaned) contig assembly using `bwa-mem2` (v2.2.1; Vasimuddin *et al*., 2019), both programs being run with default options. PCR duplicates were further removed using `rmdup` from the samtools software (v1.19.2; Li *et al*., 2009) and the realized coverage was estimated for each and every contig on the resulting `bam` file using `mosdepth` (v0.3.6; Pedersen and Quinlan, 2017) with option -Q 20 and discarding highly covered positions (*>* 300 reads corresponding to the 99th percentile of the overall coverage distribution). The mean coverage was found equal to 139X over all autosomal contigs; 82.5X over the Z–linked contigs; and 57.2X over the 17 contigs assigned to the W. Note that W–linked contigs displayed the highest proportion of highly covered positions. This clearly suggested that the two larvae used for the assembly were ZW females, i.e., the ONT data contained information to assemble the W chromosome (although at a sub-optimal coverage compared to autosomes and the Z).

### Comparative genomics

Synteny between the PPM genome and the genomes of the Lepidoptera species *Bombyx mori*, *Dry-monia ruficornis*, *Notodonta ziczac*, *Pheosia tremula*, *Ptilodon capucinus*, and *Spodoptera frugiperda*, extracted from NCBI (see Supplementary Table S6) were inspected with `GENESPACE` (Lovell *et al*., 2022) using the gene positions derived from the available GFF files, and their corresponding or-thogroups predicted with `OrthoFinder` (v2.5.5; Emms and Kelly, 2019). The phylogenetic tree was also derived from the species tree obtained with this latter program.

### Linkage Map construction

In order to estimate recombination rates and validate the scaffolding of the assembly, we aimed to construct a linkage map from genotying data for 96 offspring derived from two different batches of eggs produced after mating multiple males and females from the summer population (SP) of the Mata Nacional de Leiria (Portugal), following a protocol similar to that described in Branco *et al*. (2017). Each batch produced 48 larvae and was assumed to represent a full-sib family, with each egg batch likely originating from a single female mated with a single male, based on the known mating behavior of the PPM. Although this approach was technically the most convenient, avoiding the need to handle urticating larvae from later stages, it precluded prior knowledge of the larval sex and prevented direct genotyping of the parents.

To produce genotyping data, a Restriction-site Associated DNA (RAD) library was constructed for 96 individuals, following the protocols described in Ogden *et al*. (2013) and Gautier *et al*. (2013). Briefly, the genomic DNA was digested using the *SbfI* restriction enzyme, followed by a shearing step targeting fragments of 300–600 nucleotides. Dual indexing was used to label individual samples, with index sequences located at both fragment ends. To increase barcode diversity, each individual was assigned two distinct index combinations by varying the second-read barcode, resulting in 192 unique barcode combinations across the dataset. The resulting library was sequenced at the MGX platform on an Illumina HiSeq 2500 in paired-end mode (2×125 bp) on a single flow-cell lane. Raw sequencing data were then processed using the `process_radtags` module from the `Stacks` software package (v1.35; Catchen *et al*., 2013) to i) filter out low-quality reads; ii) verify the presence of the *SbfI* recognition site; and iii) demultiplex reads. A total of 183,087,590 read pairs (93.53% of all reads) exhibited the expected structure (i.e., a valid barcode immediately followed by the *SbfI* cut site) and passed quality filters for both reads. These represented from 899,238 to 3,129,661 (1,907,162 on average) read pairs for the different individuals (deposited on the NCBI SRA repository under the PRJNA1227736 accession ID), which were aligned onto the `Tpit_2.1` assembly using `bwa-mem2` (v2.2.1; Vasimuddin *et al*., 2019) run with default options.

Variant calling was then performed jointly on the 96 *bam* files using the multi-allelic caller imple-mented in the `bcftools` software package (v1.20; Danecek *et al*., 2021) by first computing genotype likelihoods using `mpileup` (with options -q 20 -Q 20) and directly streaming the output into the call program (with options `-mv -a GQ,GP`). Variants were subsequently filtered using `filter` from the `samtools` software (v1.19.2; Li *et al*., 2009) to keep only bi-allelic variants located more than 5 bp (option -g 5) away from any indel and with a variant quality *>*100; a read depth *>*250 across all samples; at least 90% of samples covered by at least one read; and a MAF*>*0.1. The resulting 12,107 SNPs and 1,546 indels could be assigned to 2,735 different RAD loci (with a median of 4 variants per locus). These represented 2,100 of the 6,309 that could be predicted by *in silico* digestion of the `Tpit_2.1` assembly (i.e., less than 1 kb from a *SbfI* recognition site), and 635 additional RAD sites identified by clustering variants less than 2 kb apart (i.e. +/-1 kb from a presumed restriction site) on the assembly. As a preliminary analysis, we first confirmed that the offspring comprised two full-sib families of 48 individuals each by estimating pairwise relatedness with option `--relatedness2` of `VCFtools` (v0.1.16 Danecek *et al*., 2011), excluding markers mapping to the Z chromosome and filtering genotyping data with options `--minQ 100, --minGQ 20, --minDP 5, --maxDP 250`, and `--max-missing 0.8`. Second, we sought to sex all offspring from genotyping data available for 757 SNPs (673 autosomal and 84 Z-linked) with a call rate *>*95% using the program DetSex (Gautier, 2014) run with default options and assuming both unknown individual sex and SNP chromosome type. Linkage map construction was carried out with the Lep-MAP3 software package (Rastas, 2017), which is optimized for large datasets and supports the use of genotype likelihood (GL) data. As our primary goal was to estimate recombination rates per unit of physical distance, we first assumed that i) the physical map order is correct; and ii) meioses from the female parent were achiasmatic (Wang, 2011). Briefly, the `ParentCall2` module was used to infer parental genotypes from the offspring GL and remove non-informative SNP (with option `removeNonInformative=1`). The resulting data set was further filtered using the `Filtering2` to discard SNP displaying high segregation distortion (`dataTolerance=0.01`) and excess number of missing genotypes (`missingLimit=2`). To maximize the informativeness of each RAD locus and reduce redundancy, closely linked SNPs (within 2 kb) were collapsed into a single representative marker. This strategy also mitigated complications in linkage map construction that can arise when SNPs within the same RAD tag are informative in different families (e.g., inconsistent marker ordering across pedigrees). Specifically, a custom awk script was used to select the SNP with the highest number of informative meioses within each RAD locus. In cases where all SNPs were informative for only one family, but two SNPs could be identified as informative in each of the two families, these were merged to form a single representative marker, thereby optimizing the informativeness of the collapsed RAD locus. We used the `OrderMarkers2` module to compute genetic distances and evaluate marker order for each chromosome of the physical map, setting the `recombination2=0` option to account for the assumed achiasmatic female meioses. Finally, we attempted to assign the two markers from the unplaced scaffolds to autosomes using the `SeparateChromosome2` module, testing various thresholds for the LOD score (`lodLimit` option) and recombination fraction (theta option).

### TE and Automatic Gene Annotation

Repetitive elements were annotated using `EarlGrey` (v6.0.1; Baril *et al*., 2024), with the RepeatMasker search term set to -r “`Arthropoda`”, and additional options enabled for clustering the TE library to reduce redundancy (`-c yes`), removing putative spurious TE annotations shorter than 100 bp (`-m yes`), running HELIANO for detection of Helitrons (`-e yes`), increasing to 20 the number of iterations to BLAST, Extract and Extend during the refinement process (`-i 20`), and generating a soft-masked genome (`-d yes`).

We further carried gene prediction and annotation of the genome by combining both RNA-Seq and protein evidence using `Braker3` (v3.0.7; Gabriel *et al*., 2024). Gene expression data was recovered from Dorkeld *et al*. (2023) and downloaded from NCBI (Bioproject: PRJNA663237). Briefly, this data set encompasses an overall 55.7 Gb of sequences and consists of 27 RNA-Seq libraries, each constructed from pools of individuals sampled in two Italian and two Portuguese populations for nine different developmental stages (eggs, all five larval stages, early and late pupal stages, and adults, with some stages missing for some populations; Kerdelhué, 2023). RNAseq libraries were aligned onto the unmasked assembly using `STAR` (v2.7.10b; Dobin *et al*., 2012). The resulting bam files were then used with the Arthropoda protein sequences downloaded from OrthoDB (v11; Kuznetsov *et al*., 2022) to derive hints along a reference genome softmasked with `EarlGrey` (v4.1.0; Baril *et al*., 2024), before training both `GeneMark-ETP` (Brůna *et al*., 2024) and Augustus for gene annotation (v3.4.0; Gabriel *et al*., 2024). Automatic functional annotation was carried out from predicted protein se-quences with the `Blast2GO` (v6.0; Conesa *et al*., 2005) pipeline using default parameters.

In addition to this general genome annotation, and given previously reported variation in phenology across the species’ range, including the presence of an allochronic population in Portugal (see Introduction), we specifically inspected gene models annotated by `Braker3` for known Lepidoptera circadian clock genes that may play a role in seasonal timing. To do this, we retrieved orthologous protein sequences for each circadian gene listed in Brady *et al*. (2021) from UniProt, aligned them to the *T. pityocampa* proteome generated by `Braker3` using `blastp` (v2.6.0; Camacho *et al*., 2009), and manually curated the top hits. Functional annotation was further validated using the Blast2GO (v6.0 Conesa *et al*., 2005) output to confirm the identity of candidate genes.

### Expert Gene Annotation

Olfaction and detoxification are two key functions in insects (e.g.; Leal, 2013; Després *et al*., 2007), and we annotated the Odorant Receptor (OR) gene family, along with 5 gene families involved in detoxification, namely Cytochrome P450s (CYP), carboxyl/cholinesterases (CCEs), glutathione S-transferases (GSTs), uridine diphosphate (UDP)-glucuronosyltransferases (UGTs) and ATP-binding cassette transporters (ABCs). For all gene families, annotations were conducted using established sets of Lepidoptera P450, CCE, GST, UGT and ABC transporter and OR proteins to interrogate the *T. pityocampa* genome via `tblastn` and `MiniProt` (Li, 2023) searches within the `Galaxy` plat-form (Giardine *et al*., 2005). In parallel, the results of the `HMMscan` analysis (Finn *et al*., 2011) were mined for the corresponding domains of each detoxification gene family. The protein dataset consisted of manually curated sequences annotated in the genomes of various species (detoxification: *Plutella xylostella*, *Chilo suppressalis*, *Bombyx mori*, *Manduca sexta*, *Helicoverpa armigera*, *Spodoptera littoralis*, *S. frugiperda*, *Epiphyas postvittana* and *Papilio xuthus*; OR: *H. armigera*, *S. littoralis*, *S. frugiperda*, *Agrotis ipsilon*, *Agrotis segetum*, *Athetis dissimilis*, and *Athetis lepigone*). All gene models were manually validated or corrected in `WebApollo` based on homology with other lepidopteran sequences and, where available, alignment with RNAseq data (see above). Candidate genes identified for annotation were searched against the NCBI nr database using `blastp` (Johnson *et al*., 2008) to verify annotations. Finally, all CYP protein sequences were submitted to D.R. Nelson to ensure naming conventions were respected.

## Results and Discussion

### A chromosome-level assembly of the *T. pityocampa* genome

As detailed in the Materials and Methods section, we combined (i) high-coverage long-read ONT sequencing (*>*100X) and (ii) high-coverage short-read whole-genome sequencing (*>*100X) to generate a highly contiguous (filtered) contig-level assembly (232 contigs totaling 615.7 Mb with N50=5.89 Mb) of the *T. pityocampa* genome, which was subsequently scaffolded using sequencing data from an Hi-C library. The final scaffolded assembly, referred to as `Tpit_2.1`, consisted of 165 scaffolds totaling 615.9 Mb (N50=12.2 Mb), corresponding to a decisive improvement over the previous `Tpit_1.0` by Gschloessl *et al*. (2018), that contained 68,292 scaffolds for a ten times smaller N50 of 164kb. In addition, the `Tpit_2.1` assembly size was about 15% larger than `Tpit_1.0`, and remained larger than the haploid genome size estimated from short-read k-mer spectra, which ranged from 564 to 573 Mb (Figure S1). This is likely to be attributable to the inclusion of repetitive sequences that are more effectively resolved by long-read sequencing.

Furthermore, as shown in Figure 1A, the BUSCO analysis using the `lepidoptera_odb10` lineage dataset indicated a fairly high level of completeness, with 98.8% of the 5,286 conserved single-copy orthologs identified as complete.

**Figure 1:**
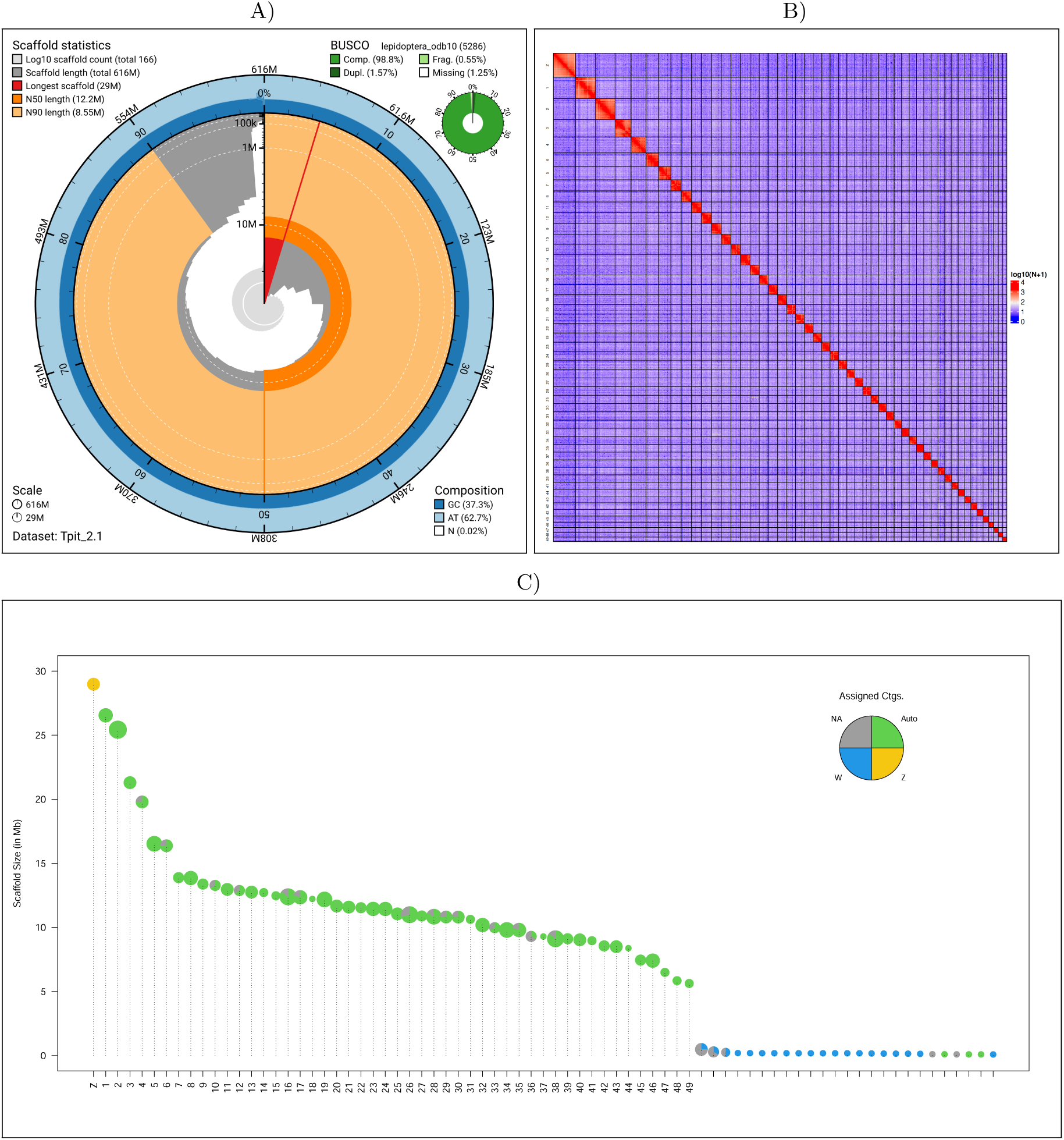
Description of the Tpit_2.1 chromosome-level assembly of the *T. pityocampa* genome. A) Snailplot built using the `BlobToolKit` (v4.3.5 Challis *et al*., 2020). B) HiC contact map at a 500 kb scale constructed from the re-mapping of HiC sequencing reads. For visualization purpose contact numbers have been rescaled (i.e., *y* = log_10_(*x* + 1) where x is the contact matrix entry). C) Identification of sex-linked and autosomal scaffolds based on the inferred chromosome type of their underlying contigs. The 75 largest scaffolds (out of the 166 ones) are ordered by decreasing sizes on the x-axis separating chromosomes, and the pie-charts represented on the y-axis give the relative proportions of contigs that are assigned to each chromosome types. The pie chart area is proportional to the number of contigs included in the scaffold.

Interestingly, the 165 scaffolds of the Tpit_2.1 assembly comprised 50 large and 115 small scaffolds (Figure 1), with a marked drop in size between the 50th (5.622 Mb) and 51st (0.472 Mb) largest scaf-folds. In addition, the 50 largest scaffolds, which we further interpreted as individual chromosomes, contained 98.8% of the assembly. As shown in Figure S4, we identified clear telomeric footprints (*>*200 repeats of the AACCT motif) at one end of 17 out of the 50 assembled chromosomes, in-cluding two chromosomes (13 and 28) with telomeric signals at both ends. These results suggest that a substantial portion of the assembly likely captures full chromosome arms, particularly among the smaller chromosomes, where telomeric motifs were frequently detected. Furthermore, 13 small scaffolds exhibited telomeric signatures, suggesting that they may represent terminal chromosome fragments that could not be integrated into the largest scaffolds. Together, these findings suggest that the assembly is not yet fully telomere-to-telomere but is nearing that stage.

Comparison of sequencing coverage between female and male allowed unambiguous assignment of the largest scaffold to the Z chromosome (Figure 1C), which was further supported by synteny analysis with several other species of Lepidoptera (Figure 2). The remaining 49 scaffolds were identified as autosomes and conventionally numbered from 1 to 49 in decreasing order of size. It should also be noted that some of the smaller scaffolds contained contigs that were assigned to the W chromosome, although these assignments should be interpreted with caution.

**Figure 2:**
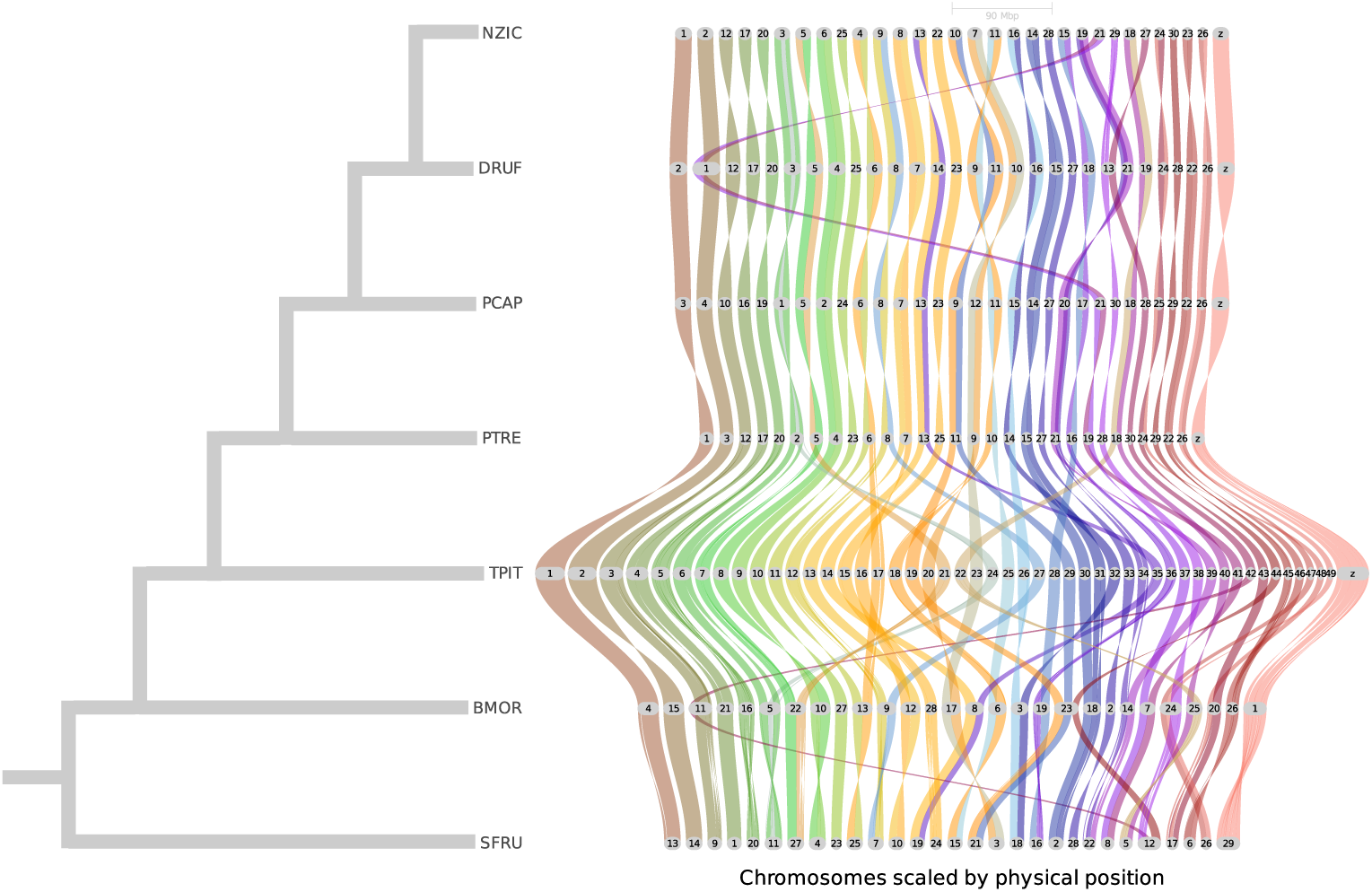
Synteny analysis between *T. pityocampa* (TPIT) and six other Lepidoptera species (BMOR: *Bombyx mori*; DRUF: *Drymonia ruficornis*; NZIC: *Notodonta ziczac*; PTRE: *Pheosia tremula*; PCAP: *Ptilodon capucinus* and SFRU: *Spodoptera frugiperda*).

Our estimate of 50 chromosomes (i.e., 49 autosomes and the Z chromosome) closely aligns with the 49 chromosomes reported for the *T. pityocampa* genome by Gschloessl *et al*. (2018) based on karyotyping, acknowledging that the high number of small chromosomes in this species made precise counting difficult. As noted previously (Gschloessl *et al*., 2018), this chromosome count lies at the upper end of the range observed in Lepidoptera, which ranges from n = 5 to n = 223, with a modal value of 31 (de Vos *et al*., 2020; Wright *et al*., 2024). Despite the higher number of chromosomes, a genome comparison revealed a high degree of conserved synteny between the *T. pityocampa* genome and some other well-sequenced Lepidoptera species (Figure 2), with evidence of chromosome fissions in *T. pityocampa*.

### Linkage Map information

From the 96 full-sibling larvae (48 per family) genotyped, a total of 46 females (20 and 26 in the two families, respectively) and 50 males were confidently identified (posterior probability *>* 0.95), consistent with an expected balanced sex ratio. Chromosome-type assignments based on linkage data were largely concordant with those inferred from the assembly. Although autosomal markers showed low genotyping errors, markers on the Z chromosome exhibited a high genotyping error rate; therefore, we restricted the construction of the linkage map to autosomes. Furthermore, the lack of parental genotypes prevented the estimation of sex-specific recombination rates, including female-specific suppression of recombination due to achiasmatic meiosis, a well-established feature in Lepidoptera (Wang, 2011). We therefore assumed that only one parent (the male) was recombining, allowing for up to 96 informative meioses in our design. Despite these limitations, after collapsing variant information within each RAD locus (see Methods and Materials section), a total of 674 markers were retained for the construction of the linkage map: 672 mapped to autosomes (ranging from 6 to 33 markers per autosome, with an average of 13.7) and two mapped to separate unplaced scaffolds. As expected given the medium marker density and relatively high recombination rate (see below), analyses based on estimated pairwise marker recombination fractions consistently produced more linkage groups than there are chromosomes in the assembly. In particular, no linkage group included markers from different chromosomes, indicating good concordance with the inferred genome structure. Of the two unplaced scaffolds, each represented by one of the 676 markers used, only one (named “U50” and 57.0 kb in length) could be integrated.

As shown in Figure S5, the estimation of genetic distances across all markers, ordered according to their physical coordinates in the Tpit_2.1 assembly and assuming no large-scale rearrangements in the SP population (an assumption supported by recent independent data, not shown), revealed a genome-wide male recombination rate of 5.06 cM/Mb. Interestingly, recombination rates varied markedly between chromosomes and, as expected, were largely explained by chromosome size: smaller chromosomes exhibited higher recombination rates, ranging from 11.6 cM/Mb on chromosome 49 to 1.98 cM/Mb on chromosome 4 (Figure S5B, Table S3). In particular, a nearly linear inverse relationship was observed between chromosome size and recombination rate for all chromosomes smaller than 15 Mb, beyond which the rate plateaued at approximately 3 cM/Mb. Comparison of the estimated genetic distances for the physical order of each of the 49 chromosomes and the optimized order estimated by the `OrderMarkers2` module of `LepMap3` (Rastas, 2017), showed that five chromosomes remained unchanged. In 35 cases, the newly proposed order led to an increase in genetic length, while in nine cases a slight decrease was observed. Overall, the cumulative genetic map length over the proposed orders increased by 14.5%, with an average change of +16.7% per chromosome (ranging from −38% to +122%). This confirms that the physical map is largely consistent with the genetic data and that attempts to improve marker order based on recombination rates do not yield convincing improvements. However, we cannot exclude the possibility that some of the proposed order changes reflect true rearrangements segregating in the population used to construct the linkage map.

Overall, the recombination rates estimated for *T. pityocampa* are consistent with previous observations in other Lepidopteran species. For example, genome-wide male recombination rates have been reported as 2.97 cM/Mb in *Bombyx mori* (Yamamoto *et al*., 2008), 180 kb/cM (equivalent to 5.56 cM/Mb) in *Heliconius melpomene* (Jiggins *et al*., 2005), 165 kb/cM (6.06 cM/Mb) in *H. erato* (Tobler *et al*., 2005), and 7.37 cM/Mb in *Leptidea sinapis*, with individual chromosome rates ranging from 3.5 to 15.3 cM/Mb (Palahí I Torres *et al*., 2023). The variation in recombination rates among chromosomes we also observed in *T. pityocampa* closely mirrors these patterns, reinforcing the generality of a size-dependent recombination landscape in holocentric genomes. This further suggests that the Tpit_2.1 assembly reflects the biologically meaningful chromosome structure.

### Automated Genome Annotation

Transposable elements (TEs) covered 49.1% of the *T. pityocampa* genome (Table S4) and were distributed relatively homogeneously across the assembly (Figure S6). Among classified elements, LINEs (Long Interspersed Nuclear Elements) retotransposons represented the largest category, comprising 12.5% of the genome, followed by DNA transposons (8.47%), rolling-circle transposons (7.97%), LTR retrotransposons (3.15%), and SINEs (Short Interspersed Nuclear Elements; 0.424%). The Penelope retroelement superfamily was detected in low abundance (0.0822%). A small portion (0.706%) was annotated as “Other” (including simple repeats, microsatellites, and RNA-derived sequences). In particular, 16.6% of the genome consisted of unclassified repetitive elements. This TE composition is broadly aligned with previous reports in Lepidoptera (Petersen *et al*., 2019; Gilbert *et al*., 2021; Perrier *et al*., 2025), although the SINE content in *T. pityocampa* appears unusually low.

More specifically, the repeat landscape of the *T. pityocampa* genome reveals distinct activity patterns across major TE classes (Figure 3). SINE elements, which represent a minor component of the repeatome, show a narrow distribution at moderate to high Kimura 2-Parameter (K2P) distances, with a peak at ∼0.3, suggesting an ancient wave of activity with little to no recent transposition. In contrast, both DNA transposons and LINE elements exhibit signatures of both ancient and recent bursts of activity. Within DNA transposons, *TcMar*, and to a lesser extent *P* elements, *Merlin*, and other *DNA* superfamilies, contribute prominently to a recent peak at low divergence (K2P *<* 0.03), whereas *hAT* and *PiggyBac* display broader distributions with peaks at higher divergence values (K2P ∼0.1), consistent with more ancient activity. Among LINEs, *L1*, *L2*, *RTE*, and *CR1* dominate in abundance and show evidence of both recent and historical activity, with *L1* and *L2* contributing most prominently to the recent burst. Finally, LTR retrotransposons, particularly *Pao* and *Gypsy*, display sharp peaks at low K2P distances (K2P *<* 0.03), indicating a relatively recent wave of transposition with limited evidence of older insertions. Overall, the landscape suggests a genome shaped primarily by recent expansions of specific DNA families (notably *TcMar*, *P* elements, and *Merlin*), LINEs (especially *L1* and *L2*), and LTR elements, with other DNA transposons contributing to older layers of TE accumulation.

**Figure 3:**
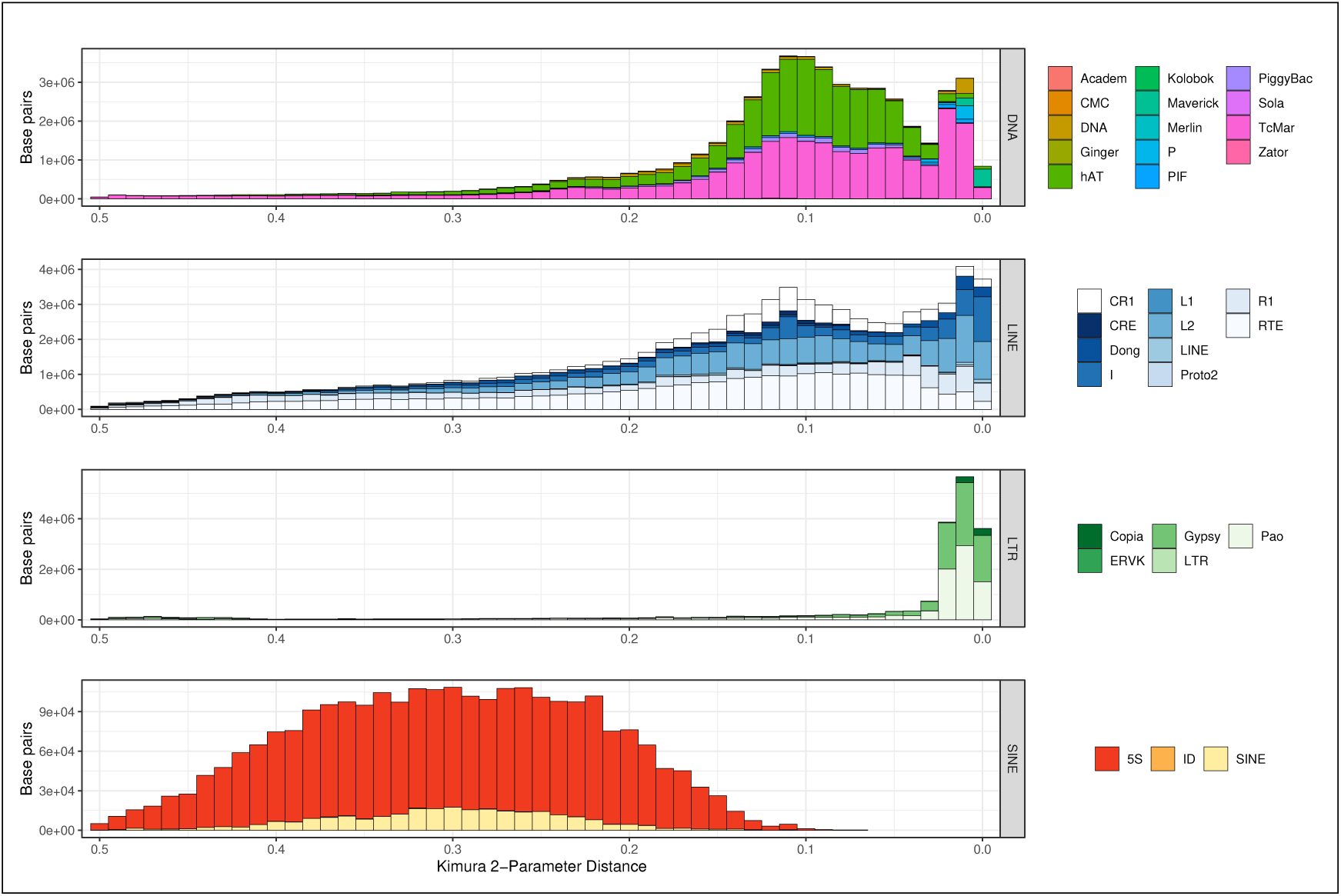
Repeat landscape of the *T. pityocampa* genome based on the annotation of the Tpit_2.1 assembly using based on EarlGrey (v6.0.1; Baril *et al*., 2024). TE are grouped grouped by super-family and plotted against sequence divergence from the consensus (K2P distance).

As summarized in Table 1, our gene annotation pipeline identified 12,898 gene models and 17,451 transcripts, which is consistent with estimates reported for other Notodontidae chromosome-level assemblies annotated using both RNA-Seq and protein evidence (min = 11,628, max = 12,888, *n* = 6; Wright *et al*., 2024). The number of genes per chromosome was positively correlated with chromosome size (Pearson’s *ρ* = 0.71) and gene density was homogeneous across the genome (Figure S6). The completeness of the *T. pityocampa* proteome was estimated at 90.7% (C:90.7% [S:89.4%, D:1.3%], F:1.9%, M:7.4%, *n* = 5,286 genes, lepidoptera_odb10) using `BUSCO` (v5.7.1; Manni *et al*., 2021) and 94.9% with OMArk (v0.3.0; Nevers *et al*., 2025). To further evaluate the quality of our protein predictions, we applied the machine learning model implemented in `PSAURON` (v1.0.1; Sommer *et al*., 2025), which yielded a score of 97.4 for *T. pityocampa* —comparable to the score obtained for the model organism *Drosophila melanogaster* (97.5).

**Table 1:** Summary statistics of the gene annotation pipeline. (Brady *et al*., 2021) (Table S5)

|  |  |
| --- | --- |
| Number of gene models | 12,898 |
| Mean gene length (bp) | 19,514 |
| Average number of exons per gene | 7.75 |
| Number of models with RNA-Seq support (%) | 12,802 (99.3) |
| Number of isoforms | 17,451 |
| Average number of isoforms per gene | 1.35 |
| Cumulative exon length (bp, %) | 28,375,678 (4.50) |
| Fraction of transcripts with functional annotation (%) | 82.5 |

Gene Ontology (GO) terms and functional annotations were assigned using `Blast2GO` for 82.5% of the 17,451 transcripts; with only 2.93% of transcripts without any significant `BLAST` hit. Among the annotated genes, we recovered 17 key loci putatively involved in the circadian clock of Lepidoptera (Brady et al., 2021) (Table S5).

### Expert Gene Annotation

We annotated 51 ORs in the *T. pityocampa* genome, out of which 37 were complete, including the obligate OR-coreceptor Orco (Larsson *et al*., 2004). This number is lower than expected for a lepidopteran species, as their genomes usually possess between 65-85 ORs (Montagné *et al*., 2020). All *T. pityocampa* ORs were found to have orthologous ORs from *S. frugiperda* and *H. armigera*, but some *S. frugiperda* and *H. armigera* OR subfamilies were devoided of any *T. pityocampa* OR, suggesting repertoire contractions. This may reflect the specialization of *T. pityocampa* on coniferous species compared to polyphagous species such as *S. frugiperda* and *H. armigera*, that may rely on a higher number of ORs to navigate to their diverse host plants, comforting the hypothesis that OR repertoire size correlates with the complexity of the species chemical ecology (Robertson, 2019). We found three TpitORs (TpitORZ-1, TpitORZ-2, and TpitORZ-3) in the group of pheromone receptors (PRs), a distinct subgroup of lepidopteran ORs, suggesting their involvement in sex pheromone communication. These three ORs localized on the Z chromosome, as observed for PRs in other species like *B. mori* (Sakurai *et al*., 2004), *Heliothis virescens* (Gould *et al*., 2010) and *Ostrinia nubilalis* (Lassance *et al*., 2011). Two of these candidate PRs (TpitORZ-2 and TpitORZ-3) presented 8 exons each and were found in close tandem in the genome, suggesting possible recent duplication. Such PRs are specialized in the detection of sex pheromone components used for intraspecific sexual communication, and are usually specifically tuned to one or a few components of the sex pheromone blend. Interestingly, the sex pheromone of *T. pityocampa* has been reported as a unique component (Quero *et al*., 1997). The discovery of three potential PRs in this species thus questions the possibility of additional, yet unknown, minor sex pheromone components in *T. pityocampa*. Alternatively, the three candidate PRs could reflect functional redundancy, as sex pheromone detection is vital for species survival. Further functional studies are required to elucidate the discrepancy between PR number and sex pheromone composition in *T. pityocampa*.

Concerning detoxification gene families, we identified a total of 236 coding genes, including 78 CYPs, 56 CCEs, 30 GSTs, 23 UGTs and 49 ABCs. This relatively moderate number of genes compared to the corresponding repertoires of other lepidopteran species (especially for UGTs and CCEs, see Supplementary Table S7) may reflect *T. pityocampa*’s narrower host plant range on coniferous species. Below we provide a summary of key findings, with detailed results and discussion available in **Supplementary Files S1-S2**.

CYP are involved in the metabolism of key endogenous substrates such as lipids and steroid hormones (Qiu *et al*., 2012; Rewitz *et al*., 2006) and are also associated to the metabolism or detoxification of xenobiotics such as plant naturals compounds and pesticides. The *T. pityocampa* genome contains 78 genes and splicing forms encoding CYPs distributed in four clans: mitochondrial (*n* = 9 genes), clan2 (*n* = 9), clan3 (*n* = 37) and clan4 (*n* = 23). Exon/intron structures were incorrectly predicted or absent from gene prediction for 25% of the CYP genes, and 58% were found in clusters, suggesting gene duplication events (Feyereisen, 2011). Interestingly, we annotated a cluster of four CYP6B on chromosome 4, with one gene CYP6B287 displaying 6 splicing forms. This CYP6B cluster was located close to the voltage-gated sodium channel protein *para*. Two new subfamilies from clan 4 were attributed to *T. pityocampa* genome, with only one member in each (CYP340DE1 and CYP341BK1).

The *T. pityocampa* genome contains 56 genes encoding putative CCEs, a multifunctional family of enzymes involved in xenobiotic detoxification, pheromone and hormone metabolism, developmental regulation, and neurogenesis. These 56 annotated genes are distributed among the 33 known CCE clades (Teese *et al*., 2010; Pearce *et al*., 2017). This total number is significantly lower than that usually observed in other Lepidoptera genomes, such as *H. armigera*, *S. frugiperda* or *B. mori*. In particular, class 1 dietary/detoxifying enzymes, known to be highly diverse among different species, comprise only 33 genes in *T. pityocampa*.

GSTs are a superfamily of multifunctional enzymes that is mainly associated with adaptation to xenobiotics, facilitating insects’ survival under chemical stresses in their environment. The *T. pity-ocampa* genome contains 30 genes encoding GSTs, which represents an intermediate number com-pared to other lepidopteran species. They are distributed across eight different classes: delta (*n* = 5 genes), epsilon (*n* = 8), microsomal (*n* = 5), omega (*n* = 4), sigma (*n* = 2), theta (*n* = 1), zeta (*n* = 3) and unknown (*n* = 2). Within the delta class GST, the GSTd2 gene showed multiple splice variants, indicating potential functional diversification.

UGTs are involved in the excretion of a wide variety of compounds including xenobiotics, endogenous metabolites, and secondary metabolites (Hung *et al*., 2019). *T. pityocampa* possesses 23 UGT genes, which is considerably fewer than other lepidopteran species (*n* = 40, *n* = 41, and *n* = 43 for *S. frugiperda*, *H. armigera* and *B. mori*, respectively). The UGT genes are distributed across eleven classes. Among those, Class 40 represents the largest expansion with seven members, suggesting particular importance for this class in *T. pityocampa* detoxification processes. Class 33 also shows significant expansion with five genes, including multiple variants of UGT33T, suggesting potential subfunctionalization or neofunctionalization. The presence of multiple gene copies within certain classes (particularly classes 33 and 40) suggests that gene duplication events may have contributed to the functional diversification of UGTs in *T. pityocampa*, and the question of whether pine hosts pose specific chemical challenges remains open (Wang *et al*., 2024).

Finally, ATP-binding cassette transporters (ABCs) use ATP hydrolysis to transport a variety of molecules such as amino acids, sugars, and even insecticides across lipid membranes. ABC trans-porters are classified into 8 subfamilies (A-H) based on similarities in their nucleotide-binding do-mains (NBD) while transmembrane domains (TMDs) are responsible for substrate specificity. The genome of *T. pityocampa* contains 49 genes encoding ABCs, which is slightly lower than observed in other lepidopteran genomes (Denecke *et al*., 2021). The largest family in *T. pityocampa* is subfamily G, with 16 members. Several genes in this subfamily are organised in pairs in the genome, with chro-mosome 13 carrying five ABCG genes. The subfamilies B and C best known for their involvement in insecticide resistance (Amezian *et al*., 2024) comprise eight and 10 members, respectively.

## Conclusion

We present a new chromosome-level genome assembly for *T. pityocampa*, which provides a highly contiguous and fully annotated reference that will be instrumental for future population genomics studies requiring chromosome-scale resolution. Compared to the previous assembly (Gschloessl *et al*., 2018), this version shows substantial improvements: it is significantly more contiguous (50 chromosome-scale scaffolds and 115 small ones, versus 68,292 contigs), slightly larger in size (615.9 Mb vs. 537 Mb), and markedly more complete, as indicated by a higher BUSCO score (98.9% vs. 83.6%). Contigs corresponding to the Z chromosome and putative W-linked regions were also iden-tified, and telomeric repeats were detected at the ends of several scaffolds, supporting near-complete chromosomal representation. This new assembly also includes both structural and functional anno-tations and we incorporated linkage map information. This resource and its annotation are already instrumental for studies on the genomic basis of both allochronic divergence (Muller *et al*., 2025) and insecticide resistance (Nouhaud *et al*., 2025) in *T. pityocampa*. Beyond its relevance for popu-lation genomics, this resource will also support comparative genomic analyses within Lepidoptera, particularly in the Notodontidae family, which encompasses more than 3,000 species, but where chromosome-level assemblies remain scarce and are so far limited to 22 species (available at NCBI as of December 2025), including five species from the distantly related Notodontinae sub-family and no representative of the Thaumetopoeinae sub-family, to which *T. pityocampa* belongs. Our anno-tated, linkage-informed assembly fills a critical gap in Lepidoptera genomics and provides a robust foundation for investigating sex chromosome evolution, local adaptation, phenology, and resistance to environmental stressors.

## Data availability statement

All raw data were deposited on the SRA repository (project PRJNA1227736). These included i) the ONT raw reads from the GridION (Run ID: SRR32477507) and PromethION (Run ID: SRR32477506) runs; ii) the PE150 Illumina short read sequences used for the polishing of the long-read assembly (Run ID: SRR32477505); iii) the HiC library paired-end sequences (Run ID: SRR32477504) used for scaffolding; iv) the PE150 WGS data generated for the two females (Run IDs: SRR32477500 and SRR32477501) and two males (Run IDs: SRR32477502 and SRR32477503) used for the identification of autosomal and sex-chromosome contigs; and v) the RAD-sequencing data for the 96 offspring of the two full-sib families (SRR33409603 to SRR33409698) used to construct linkage maps. The B29F14 BAC sequence, the *de novo* assembly and its associated annotations (including repeats, automatic gene predictions and manually curated gene models) are publicly accessible on the `LepidoDB` website https://bipaa.genouest.org/sp/thaumetopoea_pityocampa/.

## Acknowledgments

We thank Jérôme Rousselet and Patrick Pineau (URZF, INRAE Centre Val de Loire, France), for providing samples from Orléans La Source, and Susana Rocha (ISA, University of Lisbon, Portugal) for her help with SP sampling and experimental procedures in Portugal. The authors thank Muriel Latreille (UMR AGAP, INRAE, Montpellier, France) for her help with high molecular weight DNA extraction. This work was carried out in collaboration with the GeT core facility, Toulouse, France (GeT, https://doi.org/10.15454/1.5572370921303193E12), and we specially thank Roxane Boyer for support with bioinformatics analyses, Amalia Sayeh for assistance with ONT library preparation and sequencing, and Lisa Gil for assistance with Illumina short-read sequencing at the sequencing platform. MGX and GeT core facility acknowledge financial support from France Génomique National infrastructure, funded as part of “Investissement d’avenir” program managed by Agence Nationale pour la Recherche (contract ANR-10-INBS-09). We thank GenSeq technical facilities of ISEM, Montpellier, France supported by the Labex CeMEB and ANR ‘Investissements d’Avenir’ program (ANR-10-LABX-04-01), for the use of the ultrasonicator. The authors are also grateful to the Toulouse Occitanie bioinformatics platform Genotoul (Bioinfo Genotoul, https://doi.org/10.15454/1.5572369328961167E12) for providing computing resources. We finally would like to thank the DrosEU consortium for helpful discussion during stimulating meetings.

## Conflict of Interest

The authors declare no conflict of interest.

## Funding

This work was supported by the French National Agency of Research (ANR), projects GenoPheno (ANR-10-JCJC-1705-01) and LoadExp (ANR-22-CE02-0009) and by INRAE recurrent funding. The work done in Portugal was partly financed by the Fundação para a Ciência e Tecnologia, FCT-MCES, Portugal, project UID/AGR/00239/2013.

## Notes

### Competing Interest Statement

The authors have declared no competing interest.

### Summary of Updates

This version of the manuscript has been revised to include expert annotation of the odorant receptor (OR) and detoxification genes family

