## Supplementary material for "A Chromosome-Level Assembly of the Pine Processionary Moth (*Thaumetopoea pityocampa*) genome": FileS1

Supplementary file S1 for

### A chromosome-level assembly of the pine processionary moth (*Thaumetopoea pityocampa*) genome

Gautier M*, Nouhaud P, Lagnel J, Branco M, Chertemps T, Dorkeld F, François MC, Gschloessl B, Hilliou F, Jacquin-Joly E, Legeai F, Le Goff G, Lopez-Roques C, Maibeche M, Marande W, Parinello H, Sauné L, Perrier C* & Kerdelhué C*.

#### Context

Cytochrome P450s (CYP), carboxyl/cholinesterases (CCEs), glutathione S-transferases (GSTs), uridine diphosphate (UDP)-glucuronosyltransferases (UGTs) and ATP-binding cassette transporters (ABCs) are considered to be the primary families of detoxification genes in insects. Their involvement in the metabolism of both exogenous compounds, such as insecticides or allelochemicals emitted by host plant, and endogenous compounds, such as hormones and vitamins, is well established (review in Després et al. 2007). These proteins often act sequentially in three phases. First, functional groups are introduced into non-polar xenobiotics by phase I enzymes (such as CYPs or CCEs) through oxidation, hydrolysis or reduction reactions. Phase II enzymes (mainly UGTs and GSTs) conjugate these metabolites to endogenous polar compounds such as glutathione, to produce hydrophilic metabolites that can be then excreted from cells by membrane transporters (such as ABCs) during phase III.
These genes have been implicated in insecticide resistance in many insect species, either through a change in their expression levels or through a change in the catalytic activity of the proteins they encode (review in Hilliou et al., 2021). The expansion of these gene families in Lepidoptera has also often been associated with polyphagy, although monophagous species have also developed specialized detoxification pathways to metabolize the specific defenses of their hosts (Breeschoten et al., 2022). Following the three aforementioned phases, we report below the annotation of these five gene families in the pine processionary moth and compare their structural and functional diversity with those of other Lepidoptera, namely *Helicoverpa armigera*, *Spodoptera frugiperda* and *Bombyx mori*.

#### Detailed results and discussion

We identified a total of 236 detoxification encoding genes, including 78 CYPs, 56 CCEs, 30 GSTs, 23 UGTs and 49 ABCs. This is a relatively moderate number of genes compared to the corresponding repertoires of other lepidopteran species, especially for UGTs and CCEs (Table S7). This contrast is particularly pronounced with more polyphagous species like *H. armigera* and *S. frugiperda*, which encounter a broader diversity of plant secondary metabolites and thus may require larger detoxification gene repertoires. This may reflect *T. pityocampa* more specialized host plant range on coniferous species. Detailed analyses by gene families are provided below.

#### Phase I: Functionalization

**Cytochrome P450s (CYP)** genes constitute one of the largest gene families and are heme-containing monooxygenases (Werck-Reichhart, et al, 2000). CYP are involved in the metabolism of key endogenous substrates such as lipids and steroid hormones (Qiu et al., 2012; Rewitz et al., 2006) and are also associated to the metabolism or detoxification of xenobiotics such as plant naturals products and pesticides. CYPs participate in herbivory insects' abilities to overcome plant chemical defenses and are key component of successful adaptation to their host plants. A nomenclature based on sequence homology was proposed (Nebert et al., 1987, 1991). This nomenclature has since been revised, and the notion of "clan" has been proposed as a higher level of classification to account for the growing number of available sequences (Nelson, 1998). Arthropod P450s have been classified into six clans: historical clans 1, 2, 3, and 4; and mitochondrial clans 16 and 20 (Dermauw et al., 2020). *T pityocampa* genome contains 78 genes and splicing forms encoding CYPs distributed in four clans, mitochondrial, clan2, clan3 and clan4 (see Supplementary File S2, tabs 2-4 for the gene annotation details, gene sequences, and a comparison of repertoire sizes between species). CYPs names were given by D.R. Nelson. Exon/intron structures were incorrectly predicted or absent from gene prediction for 25% of the CYP genes. We annotated and named 9, 37, 23 and 9 members for clan2, clan3, clan4, and mitochondrial clan, respectively. CYP genes are often clustered in genomes, as a result of gene duplication events (Feyereisen, 2011) and 58% of the TpitCYP genes are found in cluster. We observed a cluster of four CYP6B on chromosome 4 with one gene CYP6B287 displaying also 6 splicing forms. In proximity of this CYP6B cluster we found the voltage-gated sodium channel protein *para*. Two new subfamilies from clan 4 were attributed to *T. pityocampa* genome with only one member in each, CYP340DE1 and CYP341BK1.

**Carboxyl/cholinesterases (CCEs)** are a multifunctional family of enzymes involved in xenobiotic detoxification, pheromone and hormone metabolism, developmental regulation, and neurogenesis. In insects, CCEs are divided into three main classes, subdivided into 33 clades comprising 17 functional groups (Teese et al., 2010, Pearce et al., 2017). Class 1 mainly comprises intracellular clades involved in the detoxification of food compounds and xenobiotics, class 2 mainly contains extracellular enzymes involved in the degradation of xenobiotics, pheromones and hormones, while class 3 mainly comprises non-catalytic ECEs involved in cell adhesion and neuron development. The numbers of CCEs and their assignments to clades for *H. armigera* and *B. mori* were taken from Pearce et al. (2017), and from Gouin et al. (2017) for *S. frugiperda*. *T. pityocampa* genome contains 56 genes encoding putative CCEs, which are well distributed among the different clades (see Supplementary File S2, tabs 5-7 for the gene annotation details, gene sequences, and a comparison of repertoire sizes between species). This total number is significantly lower than that usually observed in other species of Lepidoptera, such as *H. armigera*, *S. frugiperda* or *B. mori*. In particular, class 1 dietary/detoxifying enzymes, known to be highly diverse among different species, comprise only 33 genes in *T. pityocampa*. Most enzymes in this class have been associated with the detoxification of plant secondary metabolites and insecticides and therefore play an important role in the adaptation of insects to their host plants and in their resistance to insecticides (Oakeshott 2010; Pearce, Clarke et al. 2017). Several of them have also been linked to pheromone or other semiochemical processing (Durand, Carot-Sans et al. 2010). The class 2 size is within the expected range for a lepidopteran species, since 10 genes from hormone/semiochemical processing CCEs have been identified in the *T. pityocampa* genome, including two putative juvenile hormone esterases (JHE), one of which having a variant of the characteristic QSAG motif of JHE. Class 3 CCEs are generally well conserved in insects. They comprise 13 sequences in *T. pityocampa*, with clear orthologous relationships between the four lepidopteran species compared. This group includes several genes encoding carboxylesterase-like adhesion molecules (CLAMs), such as neuroligins, which are mainly associated with promoting cell adhesion during neural development, and two acetylcholinesterase genes involved in neurotransmitter hydrolysis.

#### Phase II: Conjugation

**Glutathione S-transferases (GSTs)** are a superfamily of multifunctional enzymes that is mainly associated with xenobiotic adaptation, facilitating insects' survival under chemical stresses in their environment. GSTs catalyze the nucleophilic attack of reduced glutathione (GSH) on electrophilic centers of xenobiotic compounds, including plant allelochemicals and synthetic insecticides. These enzymes play a crucial role in conferring xenobiotic adaptation through direct metabolism or sequestration of toxic compounds, and indirectly by providing protection against oxidative stress induced by xenobiotic exposure (Cai et al., 2022; Pavlidi et al., 2018; Lin et al., 2022). The *T. pityocampa* genome contains 30 genes encoding GSTs, which represents an intermediate number compared to other lepidopteran species such as *H. armigera* (47 genes), *S. frugiperda* (41 genes), and *B. mori* (25 genes, see Supplementary File S2, tabs 8-10 for the gene annotation details, gene sequences, and a comparison of repertoire sizes between species). They are distributed across eight different classes: delta (5 genes), epsilon (8 genes), microsomal (5 genes), omega (4 genes), sigma (2 genes), theta (1 gene), zeta (3 genes) and unknown (2 genes). The epsilon class represents the largest group with 8 members, which is consistent with the importance of this class in lepidopteran detoxification processes. The delta class GST includes five genes (GSTd1, GSTd2_iso1, GSTd2_iso2, GSTd2_iso3, and GSTd3), with GSTd2 showing multiple splice variants, indicating potential functional diversification. The microsomal GSTs comprise five genes (MGST1_1, MGST1_2, MGST2, MGST3, and MGST4), which are membrane-associated proteins involved in eicosanoid and glutathione metabolism (Enayati et al., 2005).

**UDP-glycosyltransferases (UGTs)** constitute the second major family of phase II conjugation enzymes. UGTs catalyze the covalent addition of UDP-glucose-derived cofactor to a wide variety of lipophilic substrates, including xenobiotics, endogenous metabolites, and secondary metabolites. This reaction increases the water solubility of compounds, facilitating their excretion and eliminating their biological activity (Hung et al., 2019). *T. pityocampa* possesses 23 UGT genes, which is considerably fewer than other lepidopteran species: *H. armigera* (41 genes), *B. mori* (43 genes), and *S. frugiperda* (40 genes, see Supplementary File S2, tabs 11-13 for the gene annotation details, gene sequences, and a comparison of repertoire sizes between species). The UGT genes are distributed across eleven classes: class 33 (5 genes), class 40 (7 genes), class 34 (1 gene), class 39 (1 gene), class 41 (1 gene), class 42 (2 genes), class 44 (1 gene), class 46 (2 genes), class 47 (1 gene), class 48 (1 gene), and class 50 (1 gene). Class 40 represents the largest expansion with seven members

(UGT40F1, UGT40L1, UGT40M1, UGT40Q1, UGT40R1, UGT40R2, and a second UGT40F1), suggesting particular importance for this class in *T. pityocampa* detoxification processes. Class 33 also shows significant expansion with five genes (UGT33J1, UGT33T1, UGT33T2, UGT33T3, and UGT33W1), including multiple variants of UGT33T, indicating potential subfunctionalization or neofunctionalization. The presence of multiple gene copies within certain classes (particularly classes 33 and 40) suggests that gene duplication events have contributed to the functional diversification of UGTs in *T. pityocampa*, potentially enabling the species to cope with the specific chemical challenges posed by its pine hosts (Wang et al., 2024).

#### Phase III: Elimination/Export

**ATP-binding cassette transporters (ABCs)** are transporters that use ATP hydrolysis to transport a variety of molecules such as amino acids, sugars, and even insecticides across lipid membranes. A functional transporter has two cytosolic nucleotide-binding domains (NBDs) that bind and hydrolyze ATP, and two transmembrane domains (TMDs). Some ABCs have all four domains (full-transporter, FT), while others are half-transporters (HT) and require assembly into homo- or heterodimers to be functional. ABC transporters are classified into subfamilies from A to H based on similarities in their NBD domains while TMDs are responsible for substrate specificity. The genome of *T. pityocampa* contains 49 genes encoding ABCs (see Supplementary File S2, tabs 14-16 for the gene annotation details, gene sequences, and a comparison of repertoire sizes between species), which is slightly lower than what is observed in other sequenced Lepidoptera, such as *H. armigera*, *S. frugiperda* and *B. mori* (Denecke et al., 2021). However, it should be noted that these lepidopterans have a similar number in subfamilies containing few genes, E, D, F and H, with two, one, three and three members respectively. Subfamilies E and F are highly conserved in insects, are characterized by the absence of TMD domains and are therefore not transporters (Dermauw and Van Leeuwen, 2014). Their function is essential and related to ribosome biogenesis and translation regulation (Andersen and Leevers, 2007; Barthelme et al., 2011), while subfamilies D and H are HTs for which the function is relatively unknown in insects. The largest family in *T. pityocampa* is subfamily G, with 16 members. Historically, this family is the oldest known among insects thanks to the pioneering work of Thomas Morgan (Morgan, 1910) on the *white* gene in *D. melanogaster*, which role is to transport pigment precursors in the developing eyes (Ewart and Howells, 1998). Several genes in this subfamily are organized in pairs in the genome, with chromosome 13 carrying five ABCG genes. Subfamily A comprises six genes in *T. pityocampa* that are FT. In mammals, ABCAs play a role in lipid transport (Albrecht and Viturro, 2007). A recent study shows that ABCAs also regulate lipid homeostasis in *D. melanogaster* (Chen et al., 2025). In particular, ABCA protein engulfment ABC transporter in the ovary (Eato) regulates the exposure to lipid signals necessary for neuronal debris clearance (Chen et al., 2025). The last two subfamilies, B and C, have eight and 10 members respectively and are the best known in insecticide resistance (Amezian et al., 2024). These subfamilies are also known as multidrug-resistance proteins (MDRs) or P-glycoproteins (p-gps) for B and multidrug-resistance associated proteins (MRPs) for C. In Lepidoptera, for example, mutations in ABCC2 confer resistance to certain toxins produced by *Bacillus thuringiensis* while overexpression of p-gps increases the efflux of insecticides (Amezian et al., 2024).
