## Supplementary Tables and Figures for "A Chromosome-Level Assembly of the Pine Processionary Moth (*Thaumetopoea pityocampa*) genome"

| | GridION Run (Q>9) | Promethion Run (Q>9) |
|---|---|---|
| **SRA accession ID** | SRR32477507 | SRR32477506 |
| **Nb. reads** | 39,972 (16,783) | 6,945,559 (5,055,546) |
| **Total yield in Gbp** | 0.513 (0.249) | 102.4 (81.54) |
| **Average read length in bp** | 12,845 (14,825) | 14,750 (16,130) |
| **N50 read length in bp)** | 21,316 (21,428) | 22,070 (22,271) |
| **Longest read in bp** | 103,053 (103,053) | 1,149,683 (157,788) |
| **Mean Q score** | 7.89 (10.1) | 10.2 (11.8) |
| **% reads with Q>10** | 21.1 (50.3) | 64.7 (88.9) |
| **Nb of active pores** | 88 (73) | 2,705 (2,684) |
| **Coverage (with GS=616 Mb)** | 0.83 (0.40) | 166.3 (132.4) |

Table S1: Summary statistics for the long reads obtained with the two ONT sequencing runs. Metrics are reported for either all reads or considering only reads with a quality score Q>9 (in parentheses).

| ID | Sex | SRA ID | Mreads (Gb) after filtering | % duplicate (`fastp`) | Mreads mapped (% prop. paired) | Realized median coverage |
|---|---|---|---|---|---|---|
| SP-M1 | M | SRR32477503 | 149.26 (18.6) | 1.35 | 132.6 (97.4) | 28 |
| SP-M2 | M | SRR32477502 | 173.82 (21.7) | 1.25 | 154.3 (97.5) | 33 |
| SP-F3 | F | SRR32477501 | 155.71 (19.5) | 1.39 | 135.2 (97.4) | 29 |
| SP-F4 | F | SRR32477500 | 156.60 (19.6) | 1.52 | 135.5 (97.3) | 29 |

Table S2: Summary statistics for the short-read sequence data generated for the two males and females used for the identification of sex-linked and autosomal scaffolds.

| Chromosome | Nb. SNPs | Physical Lengh in Mb (% covered) | Genetic Lengh in cM | cM to Mb |
|---|---|---|---|---|
| 1 | 33 | 25.8 (97.1) | 80.2 | 3.11 |
| 2 | 28 | 24.6 (96.7) | 71.9 | 2.92 |
| 3 | 25 | 19.8 (92.8) | 62.5 | 3.16 |
| 4 | 21 | 16.9 (85.5) | 33.3 | 1.98 |
| 5 | 17 | 15.0 (90.7) | 70.8 | 4.73 |
| 6 | 12 | 14.9 (91.2) | 43.8 | 2.93 |
| 7 | 15 | 9.32 (67.2) | 31.2 | 3.35 |
| 8 | 15 | 12.5 (90.6) | 50.0 | 3.98 |
| 9 | 6 | 10.5 (78.4) | 32.3 | 3.08 |
| 10 | 12 | 12.5 (94.2) | 36.5 | 2.92 |
| 11 | 14 | 12.1 (93.7) | 83.3 | 6.86 |
| 12 | 17 | 12.4 (96.6) | 45.8 | 3.68 |
| 13 | 11 | 12.3 (96.1) | 75.0 | 6.12 |
| 14 | 16 | 12.3 (96.9) | 45.8 | 3.72 |
| 15 | 13 | 11.6 (93.3) | 58.3 | 5.02 |
| 16 | 12 | 9.09 (73.4) | 57.3 | 6.30 |
| 17 | 9 | 11.2 (90.4) | 57.3 | 5.13 |
| 18 | 13 | 9.73 (79.7) | 42.7 | 4.39 |
| 19 | 18 | 11.7 (96.4) | 68.8 | 5.86 |
| 20 | 12 | 8.90 (76.3) | 52.1 | 5.85 |
| 21 | 19 | 11.2 (96.8) | 71.9 | 6.41 |
| 22 | 15 | 9.56 (82.9) | 54.2 | 5.67 |
| 23 | 15 | 10.1 (88.2) | 83.3 | 8.26 |
| 24 | 14 | 10.6 (92.5) | 53.1 | 5.02 |
| 25 | 15 | 10.5 (94.6) | 54.2 | 5.18 |
| 26 | 12 | 7.53 (68.6) | 28.1 | 3.73 |
| 27 | 7 | 7.64 (70.2) | 27.1 | 3.54 |
| 28 | 13 | 10.5 (97.3) | 38.5 | 3.66 |
| 29 | 19 | 9.88 (91.4) | 77.1 | 7.80 |
| 30 | 12 | 10.5 (97.4) | 44.8 | 4.26 |
| 31 | 15 | 9.74 (91.6) | 54.2 | 5.56 |
| 32 | 12 | 9.40 (92.3) | 45.8 | 4.88 |
| 33 | 19 | 8.76 (87.8) | 54.2 | 6.18 |
| 34 | 11 | 6.85 (69.9) | 70.8 | 10.3 |
| 35 | 9 | 7.58 (77.4) | 45.8 | 6.05 |
| 36 | 18 | 8.78 (94.5) | 53.1 | 6.05 |
| 37 | 10 | 8.75 (94.3) | 49.0 | 5.59 |
| 38 | 18 | 8.18 (89.8) | 58.3 | 7.13 |
| 39 | 9 | 5.90 (64.7) | 49.0 | 8.30 |
| 40 | 14 | 8.43 (93.4) | 39.6 | 4.69 |
| 41 | 9 | 7.58 (84.6) | 47.9 | 6.32 |
| 42 | 9 | 7.49 (87.6) | 49.0 | 6.54 |
| 43 | 9 | 6.59 (77.6) | 59.4 | 9.01 |
| 44 | 8 | 7.32 (87.4) | 35.4 | 4.84 |
| 45 | 6 | 5.72 (76.8) | 61.5 | 10.7 |
| 46 | 6 | 7.16 (96.7) | 63.5 | 8.88 |
| 47 | 9 | 5.17 (79.7) | 37.5 | 7.25 |
| 48 | 8 | 1.07 (18.3) | 8.33 | 7.81 |
| 49 | 13 | 3.77 (67.0) | 43.8 | 11.6 |
| ALL | 672 | 505.4 (87.2) | 2557.3 | 5.06 |

Table S3: Comparison between the physical and genetic maps of the autosomes. For each chromosome, the table lists the number of markers (from collapsed RAD loci; see main text), the physical length in megabases (Mb) corresponding to the distance between the first and last marker on the `Tpit_2.1` assembly (with the percentage of chromosome coverage in parentheses), the estimated genetic length in centimorgans (cM), and the ratio of genetic to physical distance (cM/Mb). The last row provides cumulative estimates across all chromosomes. Genetic distances were computed according to the order of the assembly, assuming no recombination in females, and using the Morgan (linear) mapping function (Rastas, 2017).

| **TE Classification** | **Coverage (in bp)** | **Copy Number** | **Genome Coverage (in %)** | **TE Family Count** |
|---|---|---|---|---|
| DNA | 52,179,497 | 154,290 | 8.47 | 327 |
| Rolling Circle | 49,056,372 | 166,094 | 7.97 | 75 |
| Penelope | 505,978 | 1,850 | 0.0822 | 4 |
| LINE | 77,212,988 | 237,713 | 12.5 | 296 |
| SINE | 2,608,800 | 14,533 | 0.424 | 6 |
| LTR | 19,417,146 | 18,904 | 3.15 | 149 |
| Other (Simple Repeat, Microsatellite, RNA) | 4,349,319 | 19,243 | 0.706 | 1,037 |
| Unclassified | 102,149,994 | 425,566 | 16.6 | 1,147 |
| Non-Repeat | 308,368,143 | – | 50.1 | – |

Table S4: Summary of TE content in the *T. pityocampa* genome based on `EarlGrey` (v6.0.1; Baril *et al.*, 2024) classification. Genomic coverage is expressed as a percentage of the total `Tpit_2.1` assembly (615.8 Mb).

| Gene (UniProt ID) Organism | *T. pityocampa* Gene ID (position) | BLAST align. length in bp (% id.; e-val) | Blast2GO annotation |
|---|---|---|---|
| timeless (Q3ZTQ6) *Danaus plexippus* | g268 1:11852874-11871503 | 1,274 (61.5; 0) | timeless |
| CRY-2 (Q0QWP3) *Danaus plexippus* | g3215 2:12457997-12482032 | 759 (69.2; 0) | cryptochrome-1-like |
| vrille (A0A4Y1PAQ0) *Helicoverpa armigera* | g718 10:8958265-8959359 | 367 (89.4; 0) | WASH complex subunit |
| FBXL (Q9VY46) *Drosophila melanogaster* | g1639 14:6461689-6487288 | 605 (37.7; $< 10^{-128}$) | F-box LRR-repeat protein |
| CRY-1 (O77059) *Drosophila melanogaster* | g3614 20:3381745-3398079 | 535 (55.9; 0) | cryptochrome-1 |
| clockwork (A0A8R2G768) *Bombyx mori* | g3864 21:3758674-3805850 | 448 (76.6; 0) | transcription factor cwo |
| doubletime (O76324) *Drosophila melanogaster* | g4464 23:10533610-10546016 | 299 (85.0; 0) | casein kinase I |
| jetlag (Q0E8T8) *Drosophila melanogaster* | g5130 26:9184097-9185514 | 255 (38.4; $< 10^{-61}$) | F-box/ LRR-repeat protein |
| Abl, dash (P00522) *Drosophila melanogaster* | g5445 28:2355794-2472236 | 443 (49.0; $< 10^{-129}$) | tyrosine-protein kinase Src42A |
| shaggy (J7G1D3) *Biston betularia* | g6509 31:622903-686762 | 272 (98.5; 0) | glycogen synthase kinase-3 beta |
| slimb (A0A212FH45) *Danaus plexippus* | g8084 38:2271243-2289937 | 490 (94.7; 0) | beta-TrCP |
| neurl4 (A0A1L4AAD6) *Drosophila melanogaster* | g11304 Z:14329649-14371365 | 1,133 (43.6; 0) | neuralized-like protein 4 |
| nep2 (A0A0B4K692) *Drosophila melanogaster* | g11305 Z:14376632-14445049 | 771 (54.9; 0) | neprilysin-2 |
| clock (Q6VRU6) *Antheraea pernyi* | g11362 Z:16936019-16961825 | 567 (66.0; 0) | circadian locomoter output cycles protein kaput |
| period (Q58A64) *Bombyx mori* | g11389 Z:17953375-17989958 | 968 (58.1; 0) | period |
| pdp1 (Q8SZT1) *Drosophila melanogaster* | g11506 Z:24582256-24733232 | 241 (71.0; $< 10^{-107}$) | Not annotated |
| bmal1, cycle (O61734) *Drosophila melanogaster* | g11509 Z:24775996-24876815 | 404 (49.0; $< 10^{-125}$) | cycle |

Table S5: Manual characterization of known Lepidopteran circadian genes in the *T. pityocampa* annotation. Homologous protein sequences were recovered from UniProt (within Lepidoptera when available, or using *D. melanogaster* if not) and blasted against the *T. pityocampa* proteome. Best hits are given with their genomic location and their Blast2GO annotation result.

| Organism (Genome version) | RefSeq or BIPAA ID (URL) |
|---|---|
| *Bombyx mori* (ASM3026992v2) | GCF_030269925.1 (https://www.ncbi.nlm.nih.gov/datasets/genome/GCF_030269925.1/) |
| *Drymonia ruficornis* (ilDryRufi1.1) | GCA_947859195.1 (https://www.ncbi.nlm.nih.gov/datasets/genome/GCA_947859195.1/) |
| *Notodonta ziczac* (ilNotZicz1.1) | GCA_918843915.1 (https://www.ncbi.nlm.nih.gov/datasets/genome/GCA_918843915.1/) |
| *Pheosia tremula* (ilPheTrem1.1) | GCA_905333125.1 (https://www.ncbi.nlm.nih.gov/datasets/genome/GCA_905333125.1/) |
| *Ptilodon capucinus* (ilPtiCapc1.1) | GCA_914767695.1 (https://www.ncbi.nlm.nih.gov/datasets/genome/GCA_914767695.1/) |
| *Spodoptera frugiperda* (7.1) | OGS7.1_20230705 (https://bipaa.genouest.org/sp/spodoptera_frugiperda_corn/download/spodoptera_frugiperda_corn/assembly_7.0/annotation_OGS7.1/) |

Table S6: List of the genomes used for the comparative genomics analysis

| Gene family | *T. pityocampa* | *H. armigera* | *B. mori* | *S. frugiperda* |
|---|---|---|---|---|
| **CYP** | 78 | 112 | 83 | 122 |
| **CCE** | 56 | 97 | 78 | 96 |
| **GST** | 30 | 47 | 25 | 41 |
| **UGT** | 23 | 41 | 43 | 40 |
| **ABC** | 49 | 54 | 54 | 67 |
| **Total** | 236 | 351 | 283 | 366 |

Table S7: Detoxification genes annotated in the genome of *T. pityocampa* and comparison with other Leipdoptera species.

### Supplementary Figures

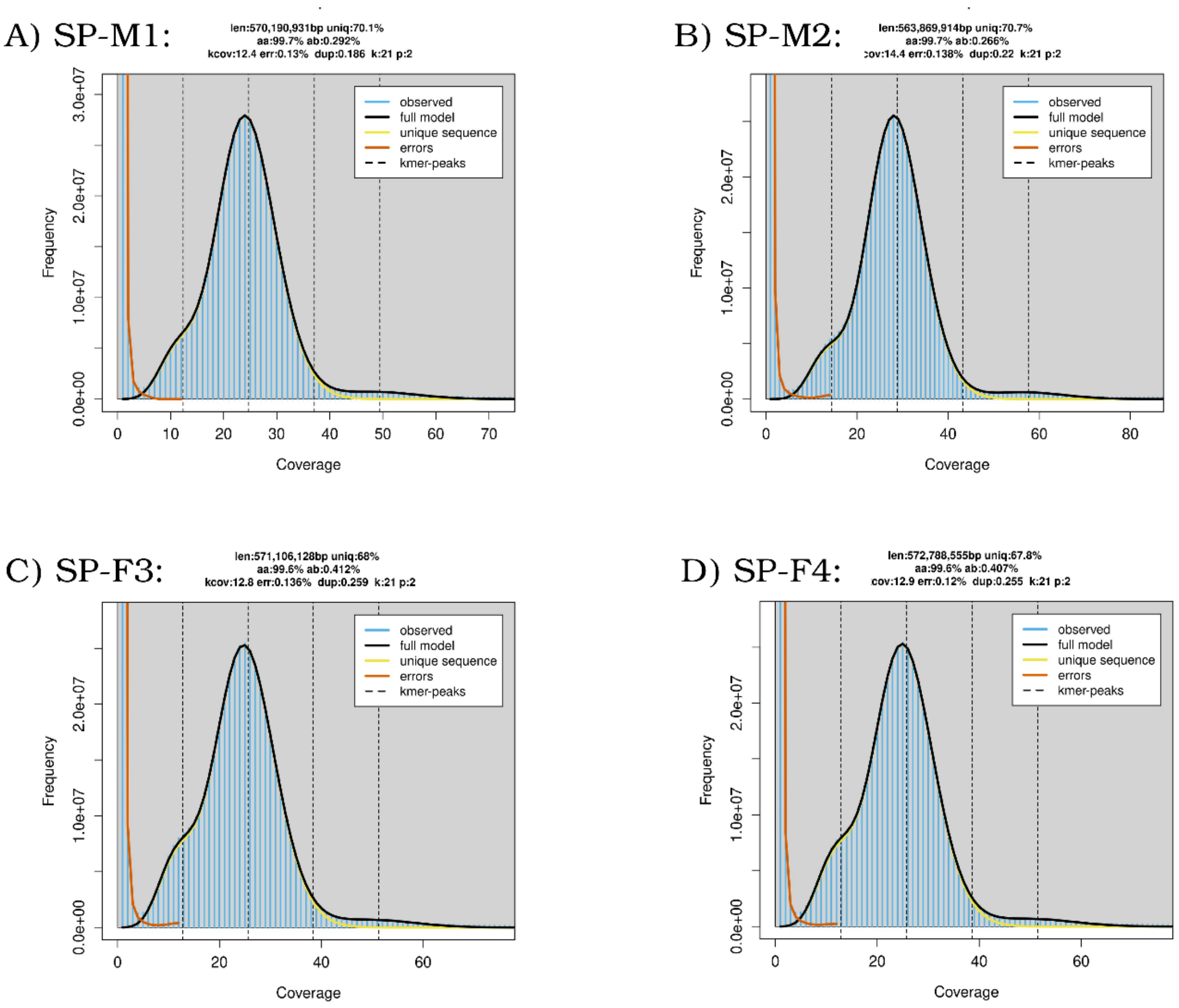


Figure S1: Estimates of the haploid genome sizes short read Whole-Genome Sequence data available for two males and two females sampled in Leiria (Portugal). The estimates were obtained using `GenomeScope` (v2.0; Ranallo-Benavidez *et al.*, 2020) from the modeling of the k-mer spectrum (with k=21) computed with `JellyFish` (Marçais and Kingsford, 2011).

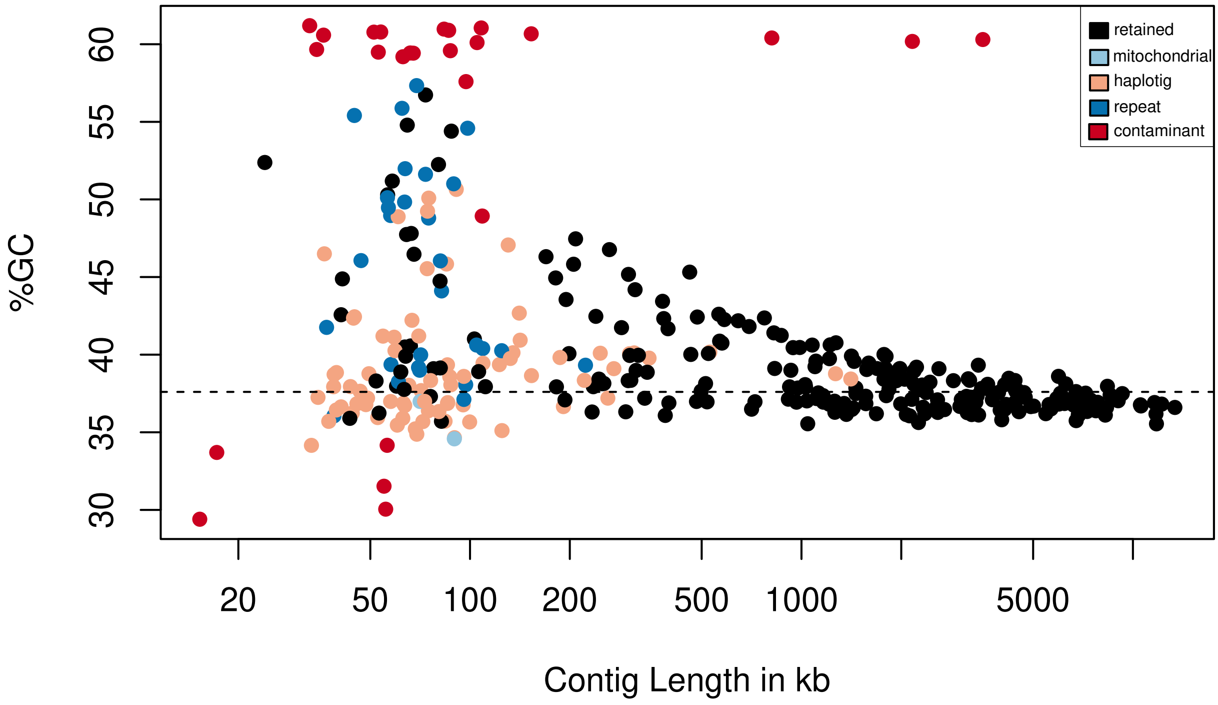


Figure S2: Distribution of the contig sizes of the polished assembly as a function of GC content. The solid line indicates the average GC content. Contigs identified as mitochondrial, contaminant or alternative chromosomal copies (see M&M section of the main text) are identified by different colors.

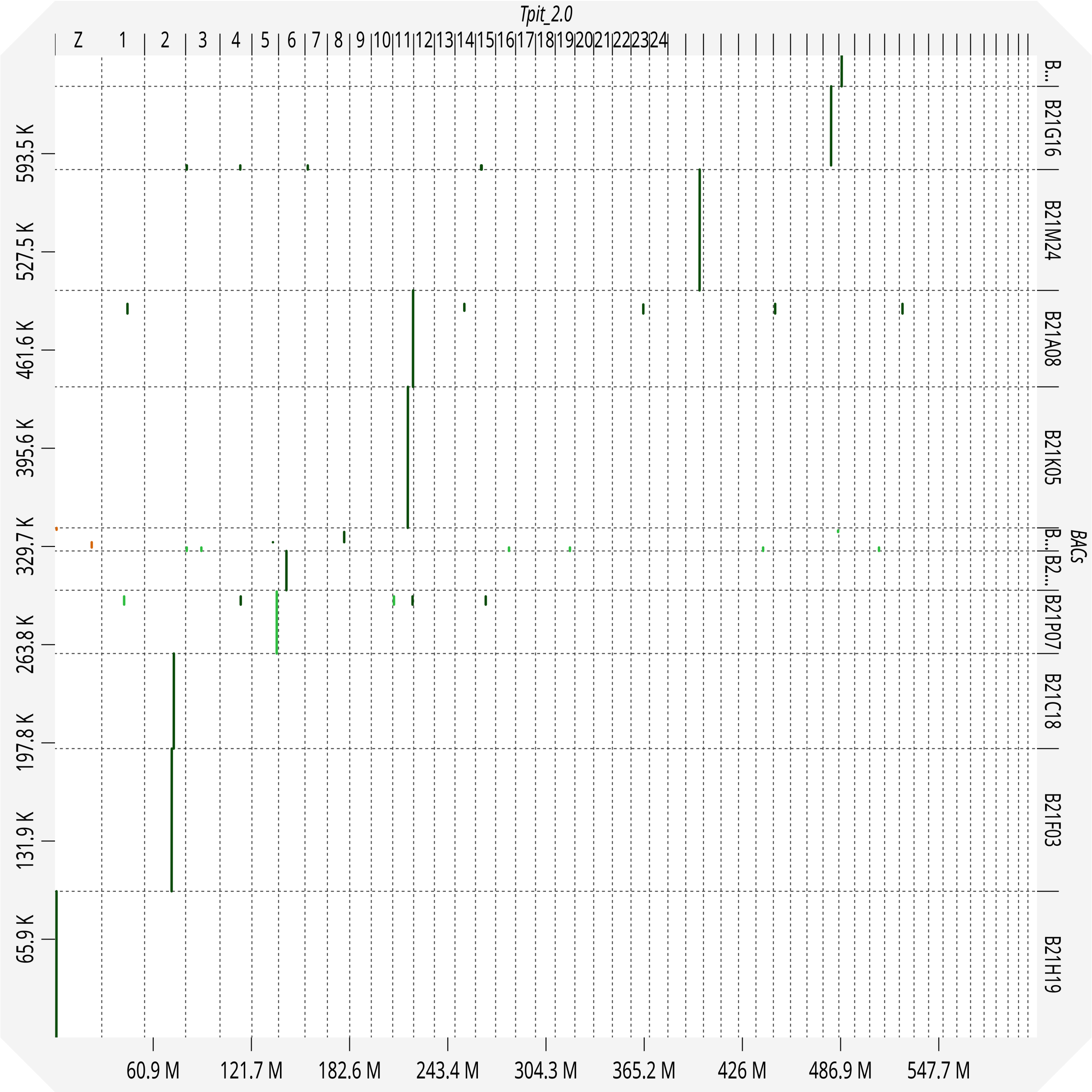


Figure S3: Dotplot generated with `dgenies` (v1.5.0; Cabanettes and Klopp, 2018) comparing the `Tpit_2.0` scaffolded assembly with the sequences of 11 BACs (Gschloessl *et al.*, 2018). Only the largest contig was considered for the 7 BACs with non contiguous sequence.

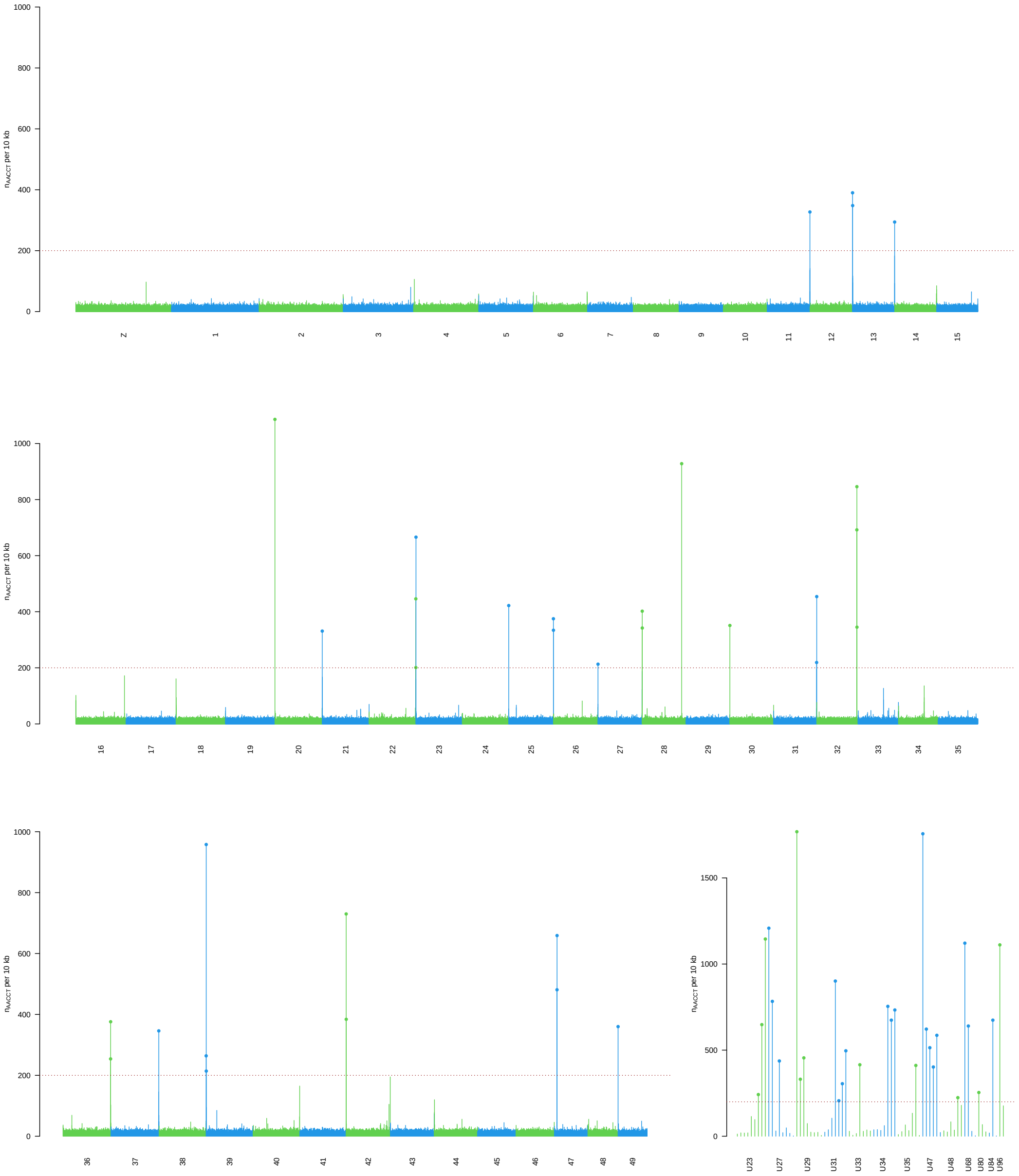


Figure S4: Detection of telomeric motif (AACCT) occurrences across assembled scaffolds. The plot shows the number of AACCT repeats per non-overlapping 10 kb window along the 50 largest chromosome-scale scaffolds (represented with alternating green and blue bars) and 13 smaller scaffolds (¡200 kb) that each contain at least one window with >200 repeats. The horizontal red dotted line indicates the arbitrary threshold of 200 repeats used to define a clear telomeric footprint. Individual windows exceeding this threshold are marked with dots.

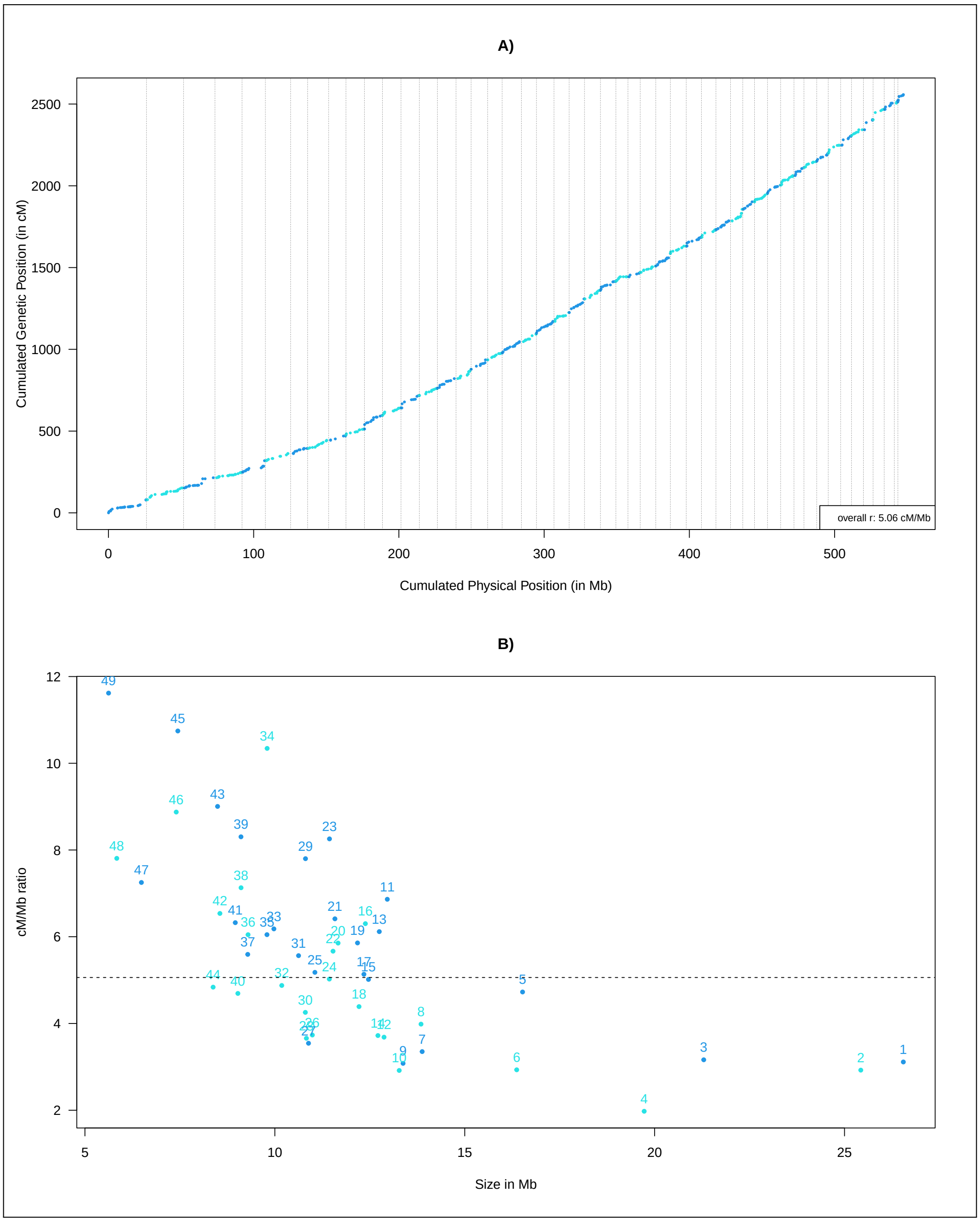


Figure S5: Comparison of the physical and genetic maps of the 49 autosomes. A) Cumulative genetic positions (in centimorgans, cM) of the 672 markers (from collapsed RAD loci; see main text and Table S3) plotted against their cumulative physical positions (in megabases, Mb) along the `Tpit_2.1` genome assembly. Chromosomes are shown in alternating colors and separated by vertical dotted lines. B) Ratio of genetic to physical distance (cM/Mb) for each chromosome as a function of its physical length on the assembly. The horizontal dotted line indicates the genome-wide average ratio. Note that genetic distances were computed according to the order of the assembly, assuming no recombination in females, and using the default Morgan (linear) mapping function as implemented in the `LepMAP3` package (Rastas, 2017).

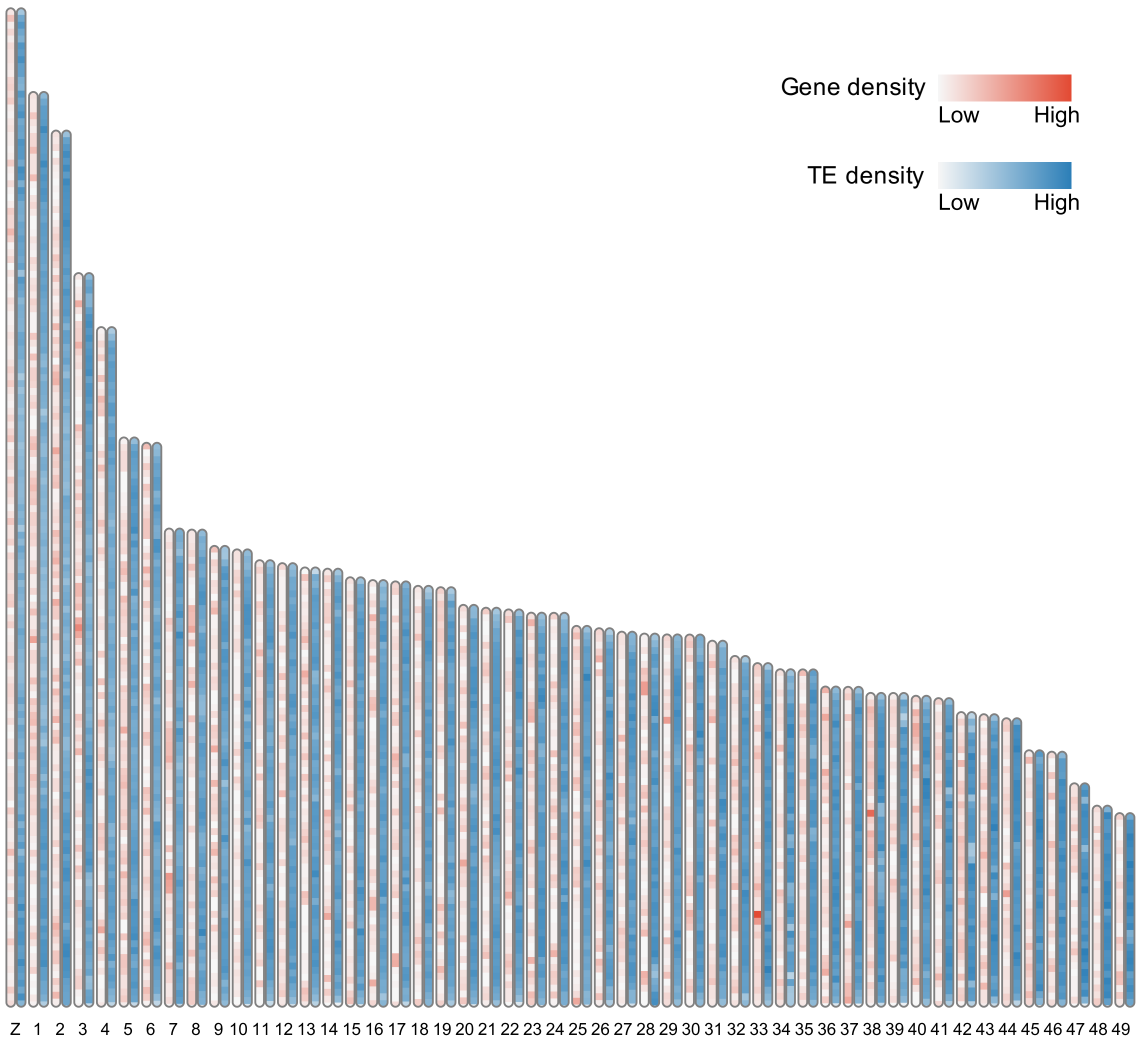


Figure S6: Gene (in red) and transposable element densities (in blue) computed over 200 kb windows along the 50 chromosomes of the *T. pityocampa* assembly.

#### Supplementary Files

**Supplementary File S1.** Detailed annotation results and discussion for detoxification genes (pdf format).

**Supplementary File S2.** Spreadsheet containing annotation results for detoxification genes (OpenDocument format).
